# DNA methylation signatures of Alzheimer’s disease neuropathology in the cortex are primarily driven by variation in non-neuronal cell-types

**DOI:** 10.1101/2022.03.15.484508

**Authors:** Gemma Shireby, Emma Dempster, Stefania Policicchio, Rebecca G Smith, Ehsan Pishva, Barry Chioza, Jonathan P Davies, Joe Burrage, Katie Lunnon, Dorothea Seiler-Vellame, Seth Love, Alan Thomas, Keeley Brookes, Kevin Morgan, Paul Francis, Eilis Hannon, Jonathan Mill

**Author notes:** Correspondence to: Professor Jonathan Mill, University of Exeter Medical School, RILD Building, Royal Devon & Exeter Hospital, Barrack Rd, Exeter. EX2 5DW. UK,.

## Abstract

Alzheimer’s disease (AD) is a chronic neurodegenerative disease characterized by the progressive accumulation of amyloid-beta and neurofibrillary tangles of tau in the neocortex. Utilizing extensive neuropathology data from the Brains for Dementia Research (BDR) cohort we performed the most systematic epigenome-wide association study (EWAS) of multiple measures of AD neuropathology yet undertaken, profiling DNA methylation in two cortical regions from 631 donors. We meta-analyzed our results with those from previous studies of DNA methylation in AD cortex (total n = 2,013 donors), identifying 334 cortical differentially methylated positions (DMPs) associated with AD pathology including methylomic variation at novel loci not previously implicated in dementia. We subsequently characterized DNA methylation in purified nuclei populations - enriched for neurons, oligodendrocytes and microglia - exploring the extent to which cortex AD-associated DMPs reflect differences manifest in specific cell populations. We find that the majority of DMPs identified in ‘bulk’ cortex tissue actually reflect DNA methylation differences occurring in non-neuronal cells, with dramatically increased effect sizes observed in microglia-enriched nuclei populations. Our study highlights the power of utilizing multiple measures of neuropathology to identify epigenetic signatures of AD and the importance of characterizing disease-associated variation in purified neural cell-types.

## INTRODUCTION

Alzheimer’s disease (AD) is a chronic and incurable neurodegenerative disease that is clinically characterized by progressive memory loss and declining cognition. Although AD is neuropathologically associated with the accumulation of extracellular amyloid-beta (Aβ) plaques and the deposit of intracellular neurofibrillary tangles of tau (NFT)^1,2^, it is also frequently accompanied by pathological features associated with other types of dementia^3,4^. Lewy-body (LB) and TDP-43 pathology, for example, are often present alongside tau and amyloid pathology in individuals with AD^4^. Despite progress in identifying both genetic^5–9^ and non-genetic risk factors for AD, the molecular mechanisms driving AD pathology remain elusive.

There is growing recognition about the importance of non-sequence-based regulatory variation in health and disease. Building on the hypothesis that epigenomic dysregulation is important in the etiology and progression of AD neuropathology^10^, we and others have identified DNA methylation differences in several regions of the brain associated with AD and also other forms of dementia including Parkinson’s disease (PD)^11–19^. A recent epigenome-wide association study (EWAS) meta-analysis, for example, reported >200 differentially methylated positions (DMPs) in the cortex associated with tau pathology^13^. There are, however, important limitations to existing studies of epigenetic variation in AD. First, because the cortex comprises a heterogeneous mix of different neural cell types - each characterized by a specific epigenetic signature - it is difficult to fully account for differences in cellular proportions between samples derived from ‘bulk’ cortex tissue. Furthermore, because the progression of AD neuropathology is associated with changes in both the number and activation of specific cell types - for example AD is associated with both the loss of neurons^20,21^ and the proliferation and activation of microglia^22,23^ – studies on bulk cortex cannot identify disease-associated variation occurring *within* individual cellular populations. Second, the clinical and neuropathological heterogeneity among patients with AD, alongside the high level of comorbidity with other types of dementia, complicates the interpretation of associations between epigenetic variation and pathology. Although existing EWAS analyses of AD have largely focused on a single pathology measure (i.e. Braak NFT staging^1,24^), the simultaneous analysis of multiple measures of different types of pathology is likely to facilitate a better understanding of the molecular mechanisms involved in disease progression.

In this study we quantified genome-wide patterns of DNA methylation in the Brains for Dementia Research (BDR) cohort, a clinically- and phenotypically-well-characterized study established with the aim of integrating standardized measures of neuropathology with detailed phenotypic and multiomic data^25^. First, we performed the most systematic EWAS of AD neuropathology yet undertaken, profiling DNA methylation across >800,000 sites in two cortical brain regions (the dorsolateral prefrontal cortex [DLPFC] and occipital cortex [OCC]) differentially impacted by AD pathology from ∼650 well-characterized donors. Second, we meta-analyzed our results with those from previous AD EWAS analyses^13^, enabling an analysis of AD-associated differential cortical DNA methylation in tissue from over 2,000 individuals. Third, we characterized genome-wide patterns of DNA methylation in purified nuclei populations enriched for neurons, oligodendrocytes and microglia from a subset of donors, exploring the extent to which AD-associated cortical differences in DNA methylation are driven by changes within specific cell populations. Our analyses identify neuropathology-associated variation at multiple novel loci not previously implicated in dementia, and show that AD-associated methylomic variation in the cortex primarily reflects differences in non-neuronal cell populations, especially microglia. This study highlights the power of utilizing multiple neuropathology measures to understand molecular pathogenesis of AD and the importance of characterizing disease-associated variation in distinct neural cell-types.

## RESULTS

### An overview of the BDR DNA methylation dataset

After stringent data pre-processing and quality control filtering (see **Methods**) the final BDR dataset comprised of DNA methylation estimates for 800,916 DNA methylation sites profiled in 1,221 tissue samples from two cortical brain regions (DLPFC and OCC) dissected from 631 donors (53% male, age range = 41-104 years, median age = 84 years, interquartile range [IQR] = 78-90 years, **Table 1**). Males were significantly younger at death compared to females (by 2.69 years, P = 2.33E-07), which is consistent with observations from epidemiological studies^26,27^. NFT pathology was quantified using Braak NFT staging^1,24^ (mean Braak score = 3.72, SD = 1.90, **Supplementary Figure S1** and **Table 1**). Amyloid pathology was quantified using both Thal phase^2^ (mean = 3.09, SD = 1.78) and neuritic plaque density scored using the CERAD classification method^28,29^ (mean = 1.69, SD = 1.28). In addition, donors were also assessed for several hallmarks of non-AD pathology including both α-synuclein pathology using Braak LB staging^30^ (mean = 1.34, SD = 2.26) and TDP-43 status (127 (22%) of 590 tested donors were TDP-43 positive).

### Alzheimer’s disease pathology is associated with altered cell-type proportions in the dorsolateral prefrontal cortex

The progression of AD pathology is associated with changes in the abundance of specific cell-types in the cortex; such changes in neural cell proportions are a major confounding factor for studies of DNA methylation and other genomic marks performed on ‘bulk’ cortical tissue^31,32^. Although several methods have been developed to derive cell-type proportion estimates from bulk DNA methylation data for use as covariates in EWAS^31,33–36^, these approaches are limited by the availability of DNA methylation reference data for specific cortical cell-types. We therefore used a fluorescence activated nuclei sorting (FANS) method recently described by our group^37^ to develop novel DNA methylation reference panels from neuron-enriched (NeuN+), oligodendrocyte-enriched (SOX10+) and microglial-enriched (NeuN-/SOX10-) nuclei populations isolated from the DLPFC from a subset of BDR donors (n = 12, see **Methods** and **Table 1**). DNA methylation profiles were generated from each purified population of nuclei using the Illumina HumanMethylation EPIC microarray and used in combination with an established algorithm^31^ to derive estimates for the proportion of each neural cell-type in individual BDR cortex samples (see **Methods**). Of note, derived relative cell proportions were significantly associated with Braak NFT stage, CERAD score and Thal phase in the DLPFC but not the OCC (see **Figure 1** and **Supplementary Table S1**), likely reflecting known differences in the progression of neuropathology across the two brain regions. In the DLPFC, increasing tau pathology was significantly associated (Bonferroni P < 0.008 [0.05/6]) with reduced neuronal cell proportion estimates (effect size = -2.74; P = 0.00011), reduced microglial proportions (effect size = -2.00; P = 0.004) and increased oligodendrocyte proportions (effect size = 1.60; P = 0.00017). This pattern was mirrored for the two measures of amyloid pathology (**Supplementary Figure S2** and **Supplementary Table S1**).

**Figure 1:**
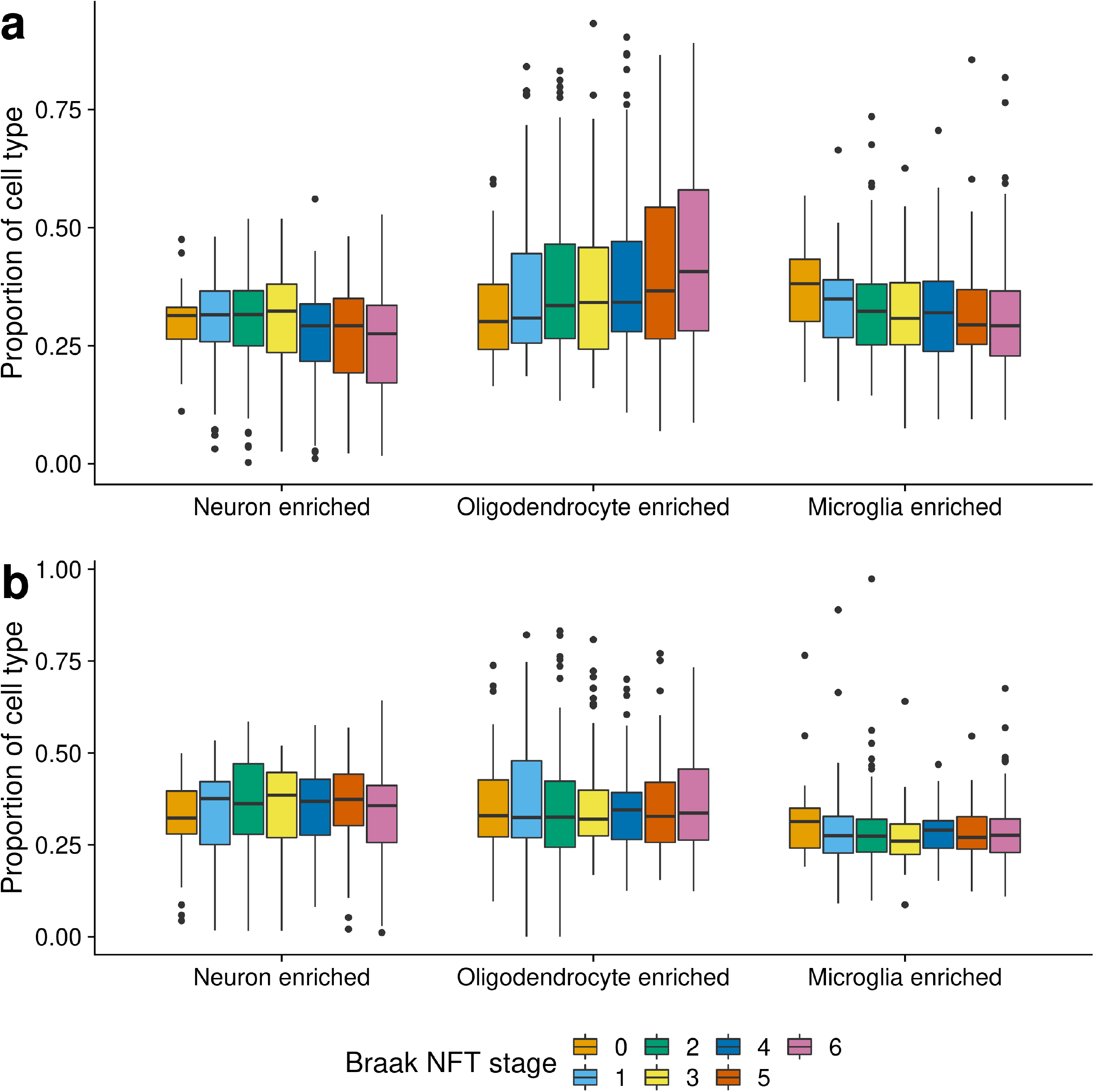
Elevated tau pathology is associated with cell proportion estimates derived from DNA methylation data in the DLPFC but not the OCC. **a)** Levels of tau pathology (measured using Braak NFT stage) are significantly associated with the proportion of neurons (effect size = -2.74, SE = 0.705, P = 1.15E-04), oligodendrocytes (effect size = 1.60, SE = 0.423, P = 1.72E-04) and microglia (effect size = -2.00, SE = 0.687, P = 0.004) in the DLPFC using neural cell proportion estimates derived from ‘bulk’ DNA methylation data. Boxplots plots for the estimated proportion of cell type across Braak NFT stages are shown, where the box in the middle represents the interquartile range (IQR), and the whisker lines represent the minimum (quartile 1 – 1.5 x IQR) and the maximum (quartile 3 + 1.5 x IQR). Tau pathology (Braak NFT stage) is shown on the x-axis split by cell-type and estimated cell proportions are shown on the y-axis. **b)** In contrast no associations between levels of tau pathology and cell proportion estimates derived from bulk DNA methylation data were observed in the OCC (P > 0.008). A similar pattern of results was found for levels of amyloid pathology as shown in **Supplementary Figure S2**).

### Multiple differentially methylated positions were associated with AD neuropathology in the cortex

We used the detailed neuropathological data available for each BDR donor to identify cortical differentially methylated positions (DMPs) associated with the accumulation of both tau (measured by Braak NFT stage) and amyloid (measured by both CERAD score and Thal Phase) pathology. We first conducted an analysis of combined AD pathology incorporating all three AD pathology measures in a model including matched DLPFC and OCC DNA methylation data from individual donors that controlled for age, sex, derived cellular proportions, experimental batch and principal component (PC) 1 (see **Methods**). We identified 67 DMPs annotated to 45 genes that were associated with the overall burden of core AD neuropathology at a stringent experiment wide significance threshold (P <9E-08) (**Figure 2a** and **Supplementary Table S2**). Of note, 32 (48%) of the significant DMPs represent sites that are specific to the Illumina EPIC array and have not been assessed in previous analyses of AD cortex undertaken that have predominantly used the preceding Illumina 450K array. The top-ranked cortical DMP associated with AD pathology was cg06913337, which was significantly hypomethylated with increasing AD pathology (P = 1.27E-10, **Figure 1b** and **1c**). Of note, this site is annotated to the *ZFPM1* gene which encodes a zinc finger protein that has been previously associated with DLB^38^ and psychosis in AD^39^.

**Figure 2:**
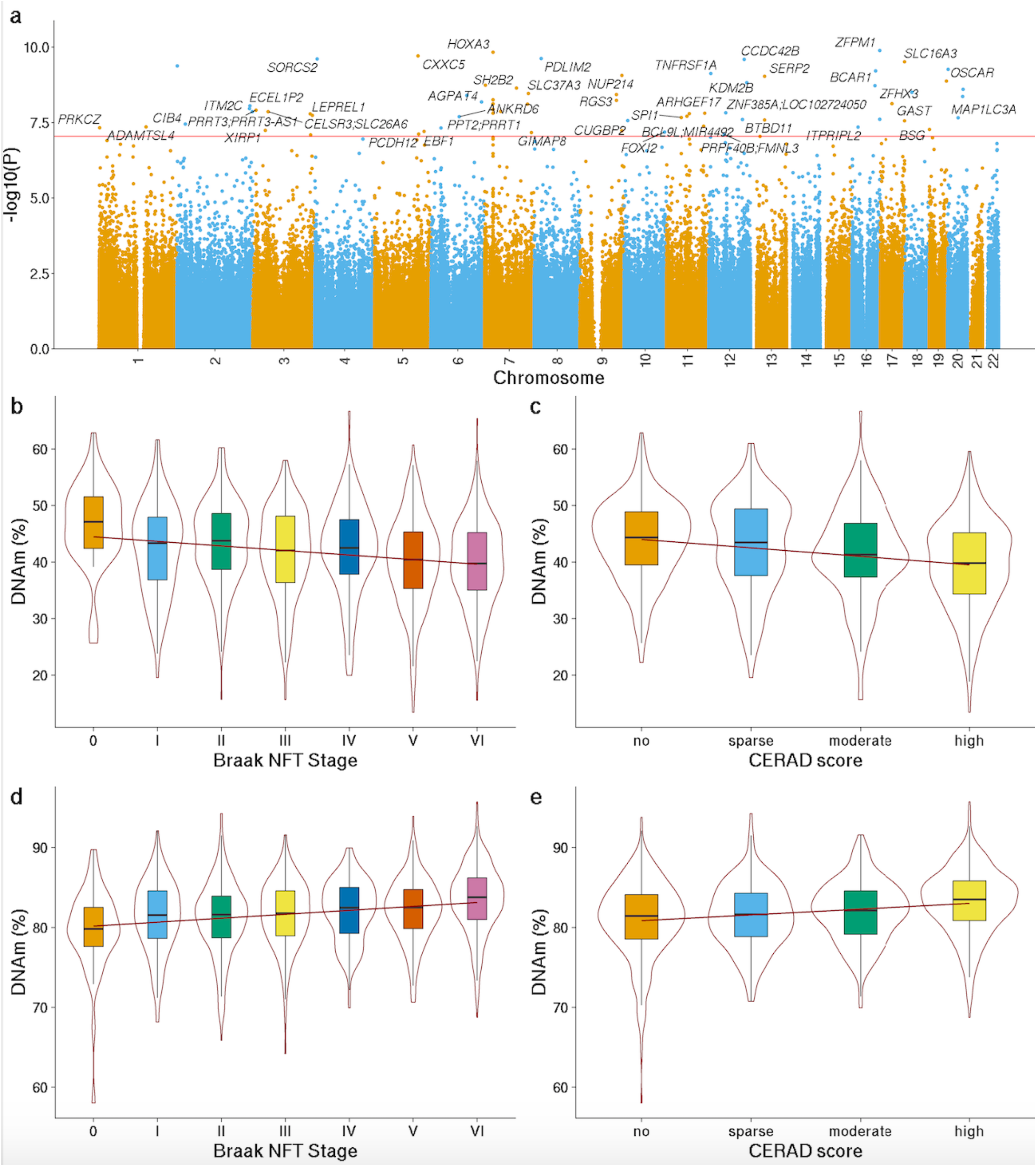
Differentially methylated positions in the cortex associated with Alzheimer’s disease neuropathology. **a)** Manhattan plot highlighting significant cortical DMPs associated with AD neuropathology (Braak NFT Stage, CERAD score, Thal phase). In total 67 DMPs associated with AD neuropathology were identified at an experiment-wide significance level (P < 9E-08). Genes annotated to significant DMPs are labelled. The x-axis depicts chromosomes 1-22 and the y-axis gives the significance level (-log10(P)) for each DNA methylation site tested. The horizontal red line represents the experiment-wide significance level (P < 9E-08). A complete list of results is given in **Supplementary Table S3** and Manhattan plots showing results from EWAS analyses of individual AD neuropathology measures are given in **Supplementary Figures S3, S6 and S9**. The top-ranked hypomethylated cortical DMP associated with AD neuropathology is cg06913337 (annotated to *ZFPM1*). Lower DNA methylation at this site is significantly associated with **b)** tau pathology (Braak NFT stage: effect size = -0.656%, SE=0.0881%, P = 2.68E-09) and **c)** amyloid pathology (CERAD score: effect size = -0.937%, SE = 0.162%, P = 6.64E-09). The top-ranked hypermethylated cortical DMP associated with AD neuropathology is cg18032191 (annotated to *TNFRSF1A*). Higher DNA methylation at this site is significantly associated with **d)** tau pathology (Braak NFT stage: effect size =0.322%, SE = 0.0598%, P = 7.20E-08) and **e)** amyloid pathology (CERAD score: effect size = 0.46%, SE = 0.0893%, P = 2.53E-07). Shown are violin plots depicting DNA methylation values (adjusted for covariates, see **Methods**) across pathology groups, where the box in the middle represents the interquartile range (IQR), and the whisker lines represent the minimum (quartile 1 – 1.5 x IQR) and the maximum (quartile 3 + 1.5 x IQR).

### Differential methylation was associated with specific Tau and Amyloid pathology measures

We next undertook analyses to identify variable DNA methylation associated with each of the three individual AD pathology measures (Braak NFT stage, CERAD score, and Thal phase). First, we identified 26 DMPs annotated to 21 genes associated with tau pathology at an experiment-wide significance threshold (P < 9E-08) (**Supplementary Figure S3** and **Supplementary Table S3**). 23 (88%) of these DMPs overlapped with sites identified in the AD neuropathology analysis. The average magnitude of effect per Braak NFT stage across these DMPs was 0.44% (SD=0.17%), with a cumulative mean DNA methylation change of 2.63% (SD=1.04%) from Braak stage 0 to VI. Of note, 22 (83%) of the DMPs were significantly hypermethylated with higher Braak NFT stage (enrichment P = 0.000267) reflecting the enrichment of hypermethylated loci observed in previous studies of tau pathology^13,16^. The top ranked DMP (cg16021126) is annotated to *SERP2*, and was significantly hypermethylated with elevated Braak NFT stage (P = 7.48E-10, effect size = 0.29% per Braak NFT stage, **Supplementary Figure S4**). *SERP2* is dysregulated in FTDP-17 (frontotemporal dementia and Parkinsonism linked to chromosome 17) iPSC-derived neurons^40^. 16 (62%) of the 26 tau-associated DMPs identified in the BDR dataset were tested in a recent meta-analysis of tau pathology performed across sites on the Illumina 450K array^13^; effect sizes for these sites were perfectly consistent across all tau-associated DMPs (100% concordant, binomial sign test P = 1.53E-05, **Supplementary Figure S5a**). It is notable that the magnitude of DNA methylation difference was approximately 2.2-fold larger in BDR than in the tau pathology meta-analysis (mean change per Braak NFT stage =0.20% [SD =0.09 %]). 6 (38%) of the 16 overlapping DMPs reached experiment-wide significance (P < 9E-08) in the previous meta-analysis and 14 (88%) reached Bonferroni significance correcting for 16 sites (Bonferroni P = 0.00313). Likewise, of the 220 DMPs identified in the tau pathology meta-analysis, 208 are included on the Illumina EPIC array and tested in the BDR dataset. These were characterized by highly consistent effect sizes observed across both analyses (100% concordant, binomial sign test P = 5.08E-61, see **Supplementary Figure S5b**); of note, effect sizes in the BDR cohort were again larger (average ∼1.2-fold larger) than those reported in the tau pathology meta-analysis.

Second, we identified 14 DMPs annotated to 12 genes associated with CERAD score (**Supplementary Figure S6** and **Supplementary Table S3**). The average magnitude of effect for the significant DMPs per unit of CERAD score was 0.57% (SD=0.16%), with a cumulative absolute mean DNA methylation difference of 2.29% (SD=0.63%) from low to high CERAD score and again an enrichment of hypermethylated sites (10 (71%) of DMPs showing higher DNA methylation with increasing pathology). The top ranked DMP (cg13515047) is annotated to *BCAR1*, which encodes a Cas scaffolding protein that acts as a functional key regulator in the pathogenesis of AD^41^, and was significantly hypermethylated with elevated CERAD score (P = 4.96E-09, effect size = 0.44%, **Supplementary Figure S7** and **Supplementary Table S3**). Finally, we identified two experiment-wide significant DMPs associated with Thal phase, both hypermethylated with increasing pathology (**Supplementary Figure S8** and **Supplementary Table S3**). The top ranked DMP (cg11658414, unannotated to a gene) was significantly hypermethylated with elevated Thal phase (P = 9.11E-09, effect size = 0.30%, **Supplementary Figure S9**).

It is well established that the neuropathological signatures of AD are correlated and higher levels of NFTs are associated with elevated amyloid burden^42^. As expected, therefore, there was a strong positive correlation in patterns of differential DNA methylation across DMPs for the individual neuropathology measures assessed in BDR (see **Supplementary Figure S10**). Effect sizes for the 26 Braak NFT stage DMPs, for example, were highly concordant (100%, binomial sign test P = 1.39E-17 across all analyses) with effect sizes at the same DNA methylation sites in analyses of the other neuropathological measures in BDR (**Supplementary Figure S11**). Additionally, when fitting the full model controlling for all AD neuropathology measures, no DMPs remained significant (P > 9E-08) for each specific measure, indicative of common effects showing consistent differences in DNA methylation across the different measures of AD neuropathology.

### Effect sizes at DMPs associated with AD pathology are correlated with those from an analysis of Lewy body and TDP-43 pathology

Because other dementia neuropathologies are frequently present alongside tau and amyloid pathology in AD we sought to explore whether DNA methylation at AD-associated DMPs was associated with Braak LB stage and TDP-43 status, two measures of common co-pathology. The samples included in our study were characterized by limited amounts of both LB and TDP-43 pathology - the majority of donors were Braak LB Stage 0 (n = 386, 72%) and TDP-43 negative (n = 463, 78%) - and we therefore had limited power to identify novel DMPs associated with either type of pathology. Elevated TDP-43 status was associated with significant hypomethylation at a single DMP (cg06423355: P = 5.47E-08, effect size = - 2.26%). Although this site is not directly annotated to any gene, it resides ∼50kb from *STK38L* which encodes a protein kinase involved in neuronal cell division and morphology and has been identified to control axonal growth in mouse hippocampal neurons^43^. Overall, effect sizes for the 67 AD pathology DMPs were found to be highly consistent between analyses of the AD (Braak NFT stage, CERAD score, Thal phase) and non-AD (Braak LB Stage and TDP-43 status) neuropathology measures (see **Supplementary Figure S12**) suggesting consistent effects across each type of neuropathology or that these effects are driven by underlying disease (i.e. a consequence rather than directly related to neuropathology).

### AD-associated differential DNA methylation is highly consistent across DLPFC and OCC

Our initial EWAS model leveraged matched DNA methylation data from both the DLPFC and OCC for each donor to maximize power to detect cortical DMPs associated with AD pathology. As expected, pathology-associated DNA methylation differences were highly consistent between both cortical regions across the 67 DMPs identified using this cross-cortex analysis model (binomial sign test P = 6.78E-21, **Supplementary Figure S13**). Given the progressive nature of AD pathology across different areas of the cortex, however, with more severe degeneration in the DLPFC compared to OCC^1,2,24^ - as reflected in our finding of pathology-associated cell proportion changes in the DLPFC but not the OCC - it is plausible that there are brain region-specific differences in AD-associated patterns of DNA methylation. Therefore, we repeated our analysis including an interaction term for brain region, identifying no significant region-specific associations with AD pathology (P > 9E-08). We also performed an EWAS of AD pathology (including the same three measures of tau and amyloid pathology) independently in each cortical region (**Supplementary Table S4**), identifying 30 significant DMPs in the DLPFC and 8 DMPs in the OCC (**Supplementary Table S5** and **Supplementary Table S6**). Although the larger number of DMPs identified in the DLPFC is consistent with the more advanced levels of AD pathology in this brain region compared to the OCC ^1,2,24^, effect sizes were strongly concordant across regions (**Supplementary Figures S14 and S15**) with one DMP (cg18100976, annotated to *PDLIM2*) being identified in both the DLPFC and OCC. Of note, *PDLIM2* encodes a protein that suppresses anchorage-dependent growth and promotes cell migration and adhesion, and has been implicated in PD by GWAS^44,45^. The consistency of findings between DLPFC and OCC suggests that variable DNA methylation at the identified DMPs is unlikely to simply reflect a consequence of neuropathology or neural cell loss.

### A meta-analysis of data from over 2,000 donors identified over 300 cortical DMPs associated with tau pathology

We combined our BDR tau pathology EWAS results with the summary statistics from a recent analysis of tau pathology performed by our group^13^, performing a cross-cortex inverse variance weighted meta-analysis of Braak NFT stage including data for 403,763 DNA methylation sites from 2,013 donors (**Supplementary Table S7**). In total we identified 334 cortical DMPs (Bonferroni P < 1.24E-07) annotated to 171 genes (**Figure 3, Supplementary Tables S8 and S9**); of note 140 (42% of total) of these DMPs represented novel associations not previously identified in the previous meta-analysis, reflecting the elevated power achieved by including the additional data from BDR donors. The top-ranked DMP, which was characterized by increasing DNA methylation with increased tau pathology (cg07061298: P = 8.06E-18, effect size = 0.32%, **Figure 3a**) is annotated to *HOXA3*; of note, previous studies have strongly implicated differential DNA methylation across the HOXA region as being associated with AD pathology^13,46,47^, and we found that 17 (5%) of the 334 meta-analysis DMPs are annotated to this genomic region (**Supplementary Figure S16**). We also confirmed other previous AD EWAS associations, including a site annotated to *ANK1* (cg05066959; P=1.16E-13, effect size =0.41%) which has been robustly associated with AD pathology in previous EWAS studies of AD ^11,15,16^ and was characterized by elevated DNA methylation with increased tau pathology (**Figure 3b**). Interestingly, several of the identified DMPs are annotated to genes that been also been implicated in GWAS analyses of AD pathology, including cg06784824 (P = 1.71E-11, effect size = 0.21%, **Figure 3c**) annotated to *SPI1*, a gene hypothesized to regulate AD-associated genes in primary human microglia^7,48^. We performed gene ontology (GO) pathway analysis of the 171 genes annotated to the significant DMPs in the cross cortex meta-analysis using *methylGSA* (see **Methods**) identifying significant enrichment of multiple pathways including pathways related to brain development and immune and inflammatory processes (see **Supplementary Table S10**). Mounting evidence suggests the immune system plays a role in the etiology of AD and other dementias^49^; both local and peripheral inflammation is triggered by the degeneration of tissues (e.g. damaged neurons and neurites) and the deposition and highly insoluble proteins such as Aβ and NFTs^49^. We subsequently repeated the meta-analysis focussing only on DLPFC samples from 1,545 individuals, identifying 300 significant DMPs annotated to 161 genes (**Supplementary Figure S17** and **Supplementary Tables S11** and **S12**). There was considerable overlap between the results from both meta-analyses with 215 DMPs being significant in both, and the direction of effect being 100% concordant between the cross-cortex DMPs (P = 2.86E-101) and DLPFC DMPs (P = 4.91E-91) (**Supplementary Figure S18**).

**Figure 3:**
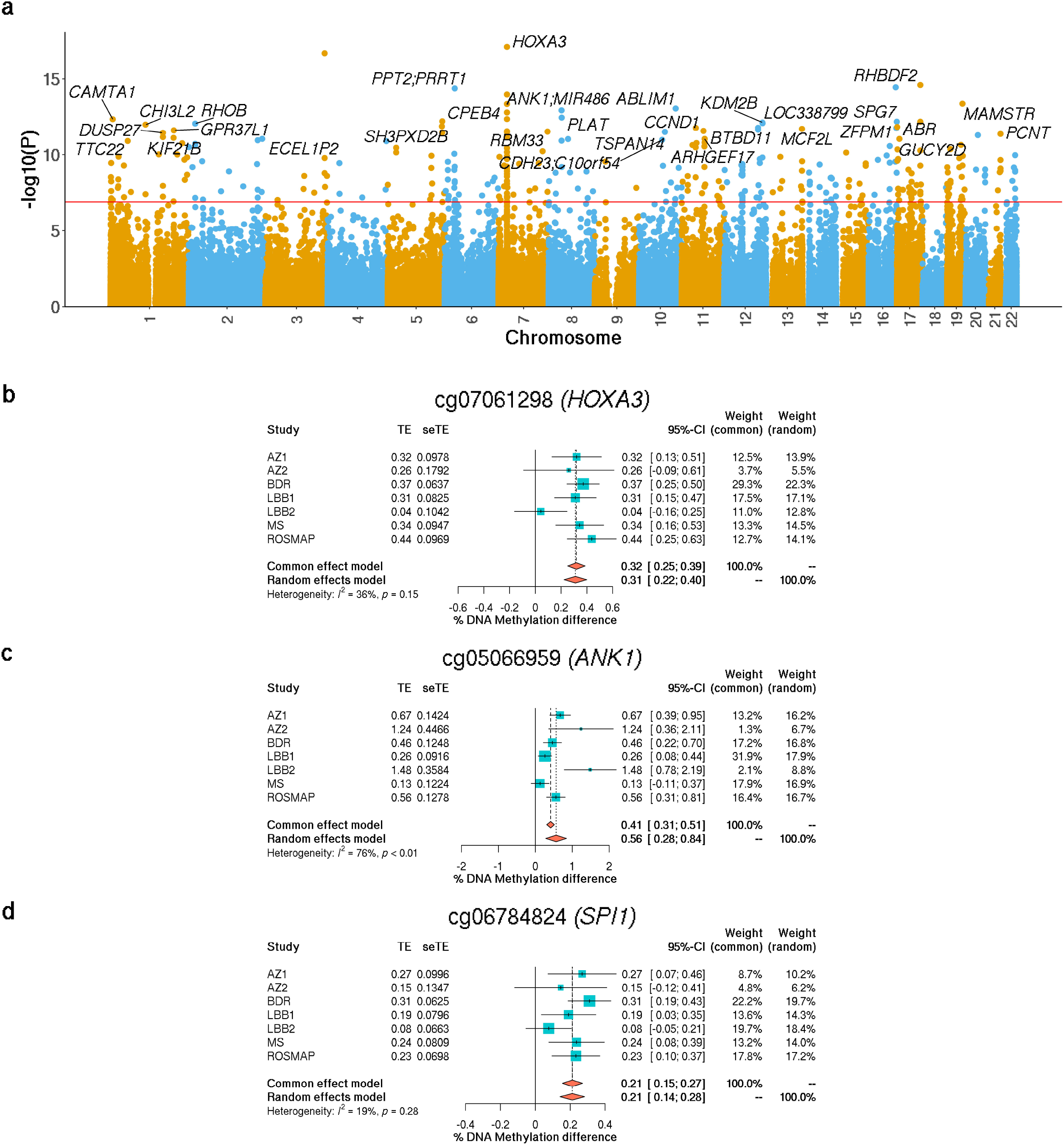
Differentially methylated positions identified in a cross-cortex meta-analysis include sites that are annotated to genes which have been previously implicated in Alzheimer’s disease. **a)** Manhattan plot highlighting significant cortical DMPs associated with Braak NFT Stage from a comprehensive EWAS meta-analysis of AD datasets (total n = 2,026 individuals). In total 334 DMPs associated with tau pathology were identified at an experiment-wide significance level (P < 9E-08). The x-axis depicts chromosomes 1-22 and the y-axis gives the significance level (-log10(P)) for each DNAm site tested. The horizontal red line represents the experiment-wide significance level (P < 9E-08). Gene annotations are given for the 50 top-ranked DMPs and a full list of results is given in **Supplementary Table S9**. Many of the DMPs associated with tau pathology have been previously implicated in AD. **b)** Elevated tau pathology is associated with **b)** hypermethylation at cg07061298 (effect size = 0.32%, SE = 0.037%, P = 8.06E-18) which is annotated to *HOXA3* that is strongly implicated in previous EWAS analyses of AD pathology, **c)** hypermethylation at cg05066959 (effect size =0.41%, SE = 0.056%, P= 1.16E-13) which is annotated to *ANK1* that is also strongly implicated in previous EWAS analyses of AD pathology, and **d)** hypermethylation at cg06784824 (effect size = 0.21%, SE = 0.032%, P= 1.71E-11) which is annotated to *SPI1* that is implicated in GWAS analyses of AD. The X-axis shows the beta effect size (% DNA methylation difference per SD increase in Braak NFT stage), with squares representing effect size and arms indicating the 95% confidence intervals.

### An analysis of purified nuclei populations shows that the majority of DMPs identified in bulk cortex tissue reflect DNA methylation differences occurring in non-neuronal cells, with dramatically increased effect sizes observed in microglia-enriched nuclei populations

Although we attempted to control for potential heterogeneity in the proportion of different neural cell-types in our analysis of bulk cortex DNA methylation by using novel reference panels generated on neuron-enriched (NeuN+), oligodendrocyte-enriched (SOX10+) and microglial-enriched (NeuN-/SOX10-) nuclei populations, our EWAS approach could not identify AD-associated differences occurring within specific cell populations. We therefore used our FANS protocol (see **Methods**) to profile DNA methylation in purified NeuN+, SOX10+ and NeuN-/SOX10-nuclei populations - in addition to a ‘total’ nuclei population reflecting the cellular makeup of bulk cortex - from DLPFC tissue from a subset of ‘low’ pathology (Braak score ≤ II, n = 15) and ‘high’ pathology (Braak score ≥ V, n = 13) donors (**Supplementary Table S13**). Of the DMPs identified in the DLPFC tau pathology EWAS meta-analysis, we obtained data for 327 sites in the purified nuclei populations (n = 327 DMPs). First we looked at between-group effect sizes in the ‘total’ nuclei population finding highly consistent DNA methylation differences to those seen in the large DLPFC meta-analysis despite the small number of samples, confirming the validity of our EWAS results (sign-test P = 7.24E-46, 87% concordant direction of effect). We then examined high vs low Braak score differences in DNA methylation at the 327 DLPFC DMPs finding a striking difference in the consistency and magnitude of effect sizes across each of the nuclei populations (**Figure 4**). Although 67 DMPs (20%) had consistent directions of effects across all nuclei populations (**Supplementary Table S14**), the NeuN-/SOX10- (microglial-enriched) population showed the most consistent between-group differences in DNA methylation (sign-test P = 1.2E-75, 96% concordant direction of effect) and was also characterized by a dramatic increase in effect sizes compared to those observed in bulk DLPFC (mean fold-change in effect size compared to bulk DLPFC =10.72, **Figure 4**). A similar pattern of differential DNA methylation was also observed in the SOX10+ (oligodendrocyte-enriched) population (sign-test P = 2.15E-10, 67% concordant direction of effect) again with an elevated effect sizes compared to bulk DLPFC, albeit to a lesser extent (mean fold-change in effect size compared to bulk DLPFC = 1.93, **Figure 4**). These results suggest that the widespread cortical DNA methylation differences associated with AD neuropathology are primarily manifest in non-neuronal cell-types, although there is evidence for pathology-associated differences in DNA methylation in neuronal cell types for a subset (5%) of DMPs (**Supplementary Table S14**).

**Figure 4:**
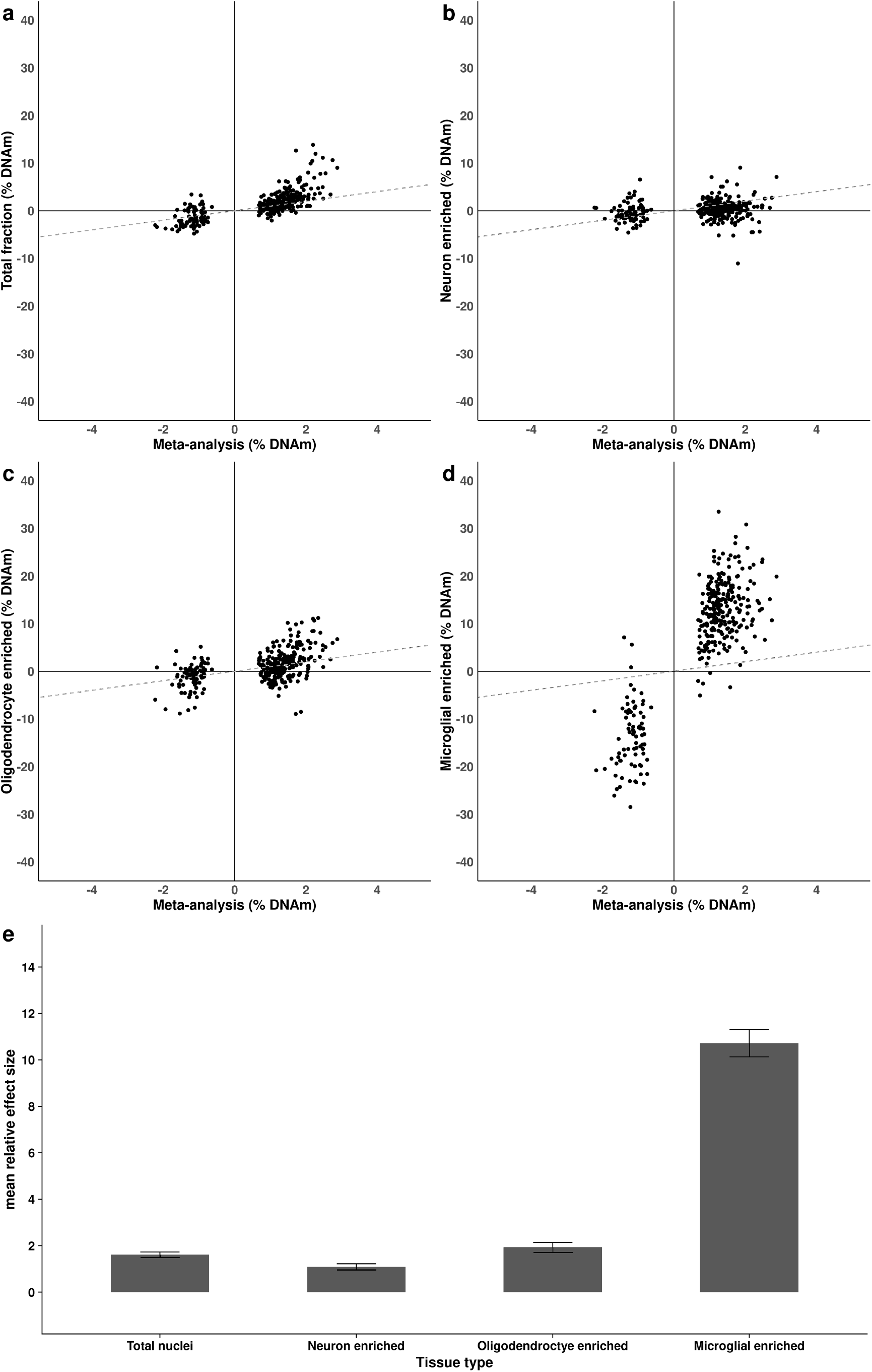
Differentially methylated positions associated with AD pathology in the cortex are largely driven by DNA methylation differences in non-neuronal cell types. We compared effect sizes for the 334 overlapping tau-associated DMPs identified in our bulk cortex meta-analysis with those at the same sites in an analysis of purified DLPFC nuclei populations from low (Braak NFT stage 0 to II) and high (Braak NFT stage > V) tau-pathology donors. Shown is a comparison of effect sizes between the meta-analysis (bulk) and the **a)** total nuclei (bulk) nuclei fraction (direction of effect = 87% concordant, sign test P = 7.24E-46); **b)** neuron enriched (direction of effect = 60% concordant, sign test P = 7.59E-05), **c)** SOX10+ (oligodendrocyte-enriched) nuclei fraction (direction of effect = 67% concordant, sign test P = 2.15E-10), and **d)** double-negative (microglial enriched) nuclei population (direction of effect = 96% concordant, sign test P = 1.2E-75). The x-axis shows effect sizes from the bulk cortex meta-analysis and the y-axis shows effect sizes for those same DMPs in each purified nuclei population. Grey dashed line represents y = x. **e)** Barplots of the mean absolute relative effect sizes in each purified nuclei population compared to the bulk cortex across the 334 tau-associated DMPs, with error bars denoting the 95% confidence intervals.

## DISCUSSION

Our study represents the most systematic analysis to date of cortical differences in DNA methylation associated with AD neuropathology. Using tissue and rich neuropathological data from 631 donors in the BDR cohort, we identified DMPs associated with levels of tau, amyloid, Lewy body and TDP-43 pathology across two cortical regions (DLPFC and OCC). We subsequently combined our results with those from previous studies of DNA methylation in AD cortex^13^, performing a meta-analysis incorporating results from over 2,000 donors and identifying 334 DMPs associated with AD pathology including many novel loci not previously identified in AD EWAS. We also characterized DNA methylation in purified DLPFC nuclei populations (enriched for neurons, oligodendrocytes and microglia) isolated from a subset of BDR donors with low and high AD pathology, exploring the extent to which pathology-associated DMPs are driven by differential DNA methylation in specific cell populations. Importantly, we find that the majority of DMPs identified in bulk cortex tissue reflect DNA methylation differences occurring in non-neuronal cells, with dramatically increased effect sizes observed in the microglia-enriched nuclei population. Our study highlights the power of utilizing multiple measures of neuropathology to identify epigenetic signatures of disease and the importance of characterizing disease-associated variation in purified neural cell-types.

Many of the pathology-associated DMPs identified in this study are annotated to genes that have previously been implicated in dementia. This includes multiple DMPs annotated to the HOXA region which has been previously identified in EWAS analyses of AD pathology^13,46,47^. The HOXA cluster is involved in the control of neuronal development, neuronal circuit organization and the regulation of post mitotic neurons^50,51^, and in addition to AD methylomic variation across the HOX region has been associated with other neurodegenerative diseases including PD, Huntington’s disease and C9ORF72-related dementia^52–54^. AD pathology-associated DMPs were also annotated to many immune related genes (e.g. *TNFRSF1A* and *OSCAR*) with gene ontology pathway analyses finding an enrichment of immune and inflammatory pathways. These findings build on existing evidence that immune dysregulation plays a key role in the etiology of AD and other dementias^49^. In addition, differential DNA methylation in the vicinity of the *SPI1* gene was identified in our cortical meta-analysis of AD pathology. *SPI1* has been identified in recent AD GWAS^7,55^ and EWAS^13^ analyses and encodes the transcription factor PU.1, a pioneer factor for myeloid macrophages and microglial populations that has been implicated in regulating genes leading to inflammatory response in AD^48,56^. This is particularly interesting in the context of our analyses of sorted nuclei populations which identified that the majority of methylomic differences associated with AD pathology occur in the microglial population.

The high overlap of DMPs and consistency of differences in DNA methylation across the different types of neuropathology assessed in BDR donors suggests that they may reflect some common signature of neurodegeneration. This could imply that these differences are a common consequence of pathology or that they reflect the known pleiotropy between different types of dementia. For example, SNPs within the HLA region, MAPT and APOE all contribute to increased risk for FTD, AD and PD^57^. Additionally, mutations in the fEOAD genes (APP, PSEN1 and PSEN2) are also present in PD cases highlighting the pleiotropic effects associated with monogenic forms of neurodegeneration^58^. Previous EWAS analyses have also identified methylomic similarities between different neurodegenerative diseases^59^ reporting significant over-representation in pathways related to brain function and immune response. The evidence for pleiotropy suggests that common pathological mechanisms likely underlie neurodegenerative disorders. Although neurodegenerative diseases differ in their neuropathological hallmarks and the specific brain regions involved, a common feature is the progressive accumulation of toxic protein deposits that ultimately lead to neuronal cell death and brain atrophy ^60^. One key strength of the BDR dataset is that multiple neuropathology measures have been collected for each individual, enabling us to identify DMPs robustly associated with overall levels of AD neuropathology and leveraging greater power than analyses based on single pathology measures. Of note, although the findings suggest there are general methylomic signatures of neuropathological burden, we cannot exclude the presence of differential DNA methylation associated with specific types of neuropathology. Interestingly the BDR effect sizes are larger than those observed in our recent meta-analysis of tau pathology^13^; this could potentially reflect cohort differences, the reduced heterogeneity in BDR, array platform differences or by the fact that association statistics for variants meeting an experiment-wide threshold tend to be overestimated^61^. In addition, the consistency in the direction of effect demonstrates how robust the EWAS results for AD pathology are across studies.

A major strength of our study is our use of FANS to purify nuclei populations from neuronal, oligodendrocyte and microglial cells on a subset of donors prior to DNA methylation profiling. This enabled us to develop a refined cell type deconvolution model that better controls for cellular heterogeneity in bulk cortex measurements of DNA methylation than previous models that only estimate the proportion of neuronal cells. Even when controlling for cell-type proportions, the bulk cortex analysis does not enable the identification of pathology-associated DNA methylation differences occurring in specific cell types. We therefore profiled DNA methylation in FANS-purified nuclei populations from individuals with high and low AD pathology to explore the extent to which differences identified in bulk tissue were driven by variation in specific cell-types. Our analyses showed that most of the DMPs identified in the bulk cortex reflect variation in non-neuronal cell-types, with the biggest effect sizes identified in nuclei from microglial cells. These results support recent work highlighting a key role for microglia in AD^62^; with the activation of microglia colocalized with amyloid plaques in the brains of individuals with AD. The larger effect sizes observed at AD-associated DMPs in the microglial-enriched population might reflect the elevated reactivity of microglia in AD compared to other neural cell types, presumably driven by cell-type-specific transcriptional signatures^62,63^.

There are several limitations that should be considered when interpreting the results of this study. First, although we attempted to control for cellular heterogeneity and profiled FANS-purified populations to compare effect sizes across different cell-types, there are some limitations to this approach – for example there is still considerable heterogeneity in each of the purified nuclei populations used to generate our deconvolution reference panels. The microglial enriched fraction, for example, will also incorporate other cell types including astrocytes, which have been implicated in neurodegeneration^64^, although co-staining of the double-negative nuclei population with the microglial marker IRF8 shows that this population is enriched for microglial cells (**Supplementary Figure 19**). Furthermore, the use of NeuN as a marker to purify neuronal nuclei is not perfect^65^. Since neurodegenerative processes are associated with atrophy of astrocytes^64^, they are an important cell-type to consider. However, it is difficult to find robust nuclear markers for this cell type. In the future a reference dataset which includes astrocytes and other cell types would be optimal. The heterogeneity of the microglial fraction may also explain the potentially surprising result of decreasing microglia proportions with increasing pathology. It is worth noting, however, that if one cell proportion decreases (e.g. the neuronal proportion) it does not necessarily mean the absolute abundance of the cell-type is changing. Despite the relatively small number of purified nuclei samples profiled in our study we were able to identify dramatically increased effect sizes in specific cell populations, highlighting the additional power gained by profiling purified cell populations.

A key limitation of epigenetic epidemiology relates to the issue of causality; it is not possible to elucidate whether the DMPs identified in this study play a causal role in driving disease pathogenesis, or whether they represent a downstream consequence of neuropathology. In this regard it is interesting that AD-associated differences identified in the OCC - a region of the cortex relatively protected from tau and amyloid pathology - were highly consistent with those identified in the DLPFC, which is affected much earlier in the disease process ^1,2,24^. This consistency across both cortical regions suggests that the AD-associated variation identified in this study do not simply represent a consequence of AD neuropathology. Of note, however, we cannot exclude the possibility that the differences identified reflect the influence of other factors related to AD pathology that were not controlled for in this study, for example environmental factors such as medication exposure.

In summary, utilizing extensive neuropathology data from the BDR cohort we have performed the most systematic EWAS of multiple measures of AD neuropathology yet undertaken. Our meta-analysis with other AD DNA methylation datasets identified 334 cortical DMPs associated with AD pathology including methylomic variation at multiple loci not previously implicated in dementia. We subsequently characterized DNA methylation in purified nuclei populations finding that the majority of DMPs identified in bulk cortex tissue reflect DNA methylation differences occurring in non-neuronal cells, with dramatically increased effect sizes observed in oligodendrocyte- and microglia-enriched nuclei populations. Our study highlights the power of utilizing multiple measures of neuropathology to understand epigenetic signatures of disease and the importance of characterizing disease-associated variation in purified neural cell-types.

## METHODS

### The Brains for Dementia Research (BDR) cohort

The Brains for Dementia research (BDR) cohort was established in 2008 and represents a network of six dementia research centers across England and Wales (based at Bristol, Cardiff, King’s College London, Manchester, Oxford and Newcastle Universities) and five brain banks (brain donations from Cardiff are banked at King’s College London)^25^. Briefly, participants >65 years of age were recruited using both national and local press (e.g. newspapers, newsletters, leaflets), TV and radio coverage as well as at memory clinics and support groups. There were no exclusion or inclusion criteria for individuals with specific diagnoses or those carrying genetic variants associated with neurodegenerative diseases; the cohort includes those with and without dementia and covers the full range of dementia diagnoses. Participants underwent a series of longitudinal cognitive and psychometric assessments and registered for brain donation.

### Post-mortem neuropathological assessment of BDR brain donations

Post-mortem brain donations to BDR undergo full neuropathological dissection, sampling and characterization by experienced neuropathologists in each of the five network brain banks using a standardized BDR protocol based on the BrainNet Europe initiative^66 67^. This protocol was used to generate a description of the regional pathology within the brain together with standardized scoring. Five variables representing four neuropathological features were used in the analyses presented in this paper: 1) Braak NFT stage which captures the progression of NFT pathology ^1,24^, 2) Thal phase which captures the regional distribution of Aβ plaques ^2^, 3) CERAD score which quantifies neuritic plaque density^29^, 4) Braak LB stage which captures the progression of α-synuclein throughout the brain^30,68^, and 5) TDP-43 status - a binary indicator of the TDP-43 inclusions, which was assessed using immunohistochemistry to identify the presence of phosphorylated TDP-43 in the amygdala, hippocampus, and adjacent temporal cortex.Braak NFT stage, Thal phase, CERAD score and Braak LB stage were analyzed as continuous variables, utilizing the semi-quantitative nature of these measures to identify dose-dependent relationships of increasing neuropathology with variable DNA methylation. TDP-43 status was analyzed as a binary variable.

### DNA methylation profiling in bulk cortex tissue

DNA methylation data were generated on two cortical regions (DLPFC and OCC) from each BDR donor. DNA was isolated from ∼100mg of tissue using the Qiagen AllPrep DNA/RNA 96 Kit (Qiagen, cat no.80311) following tissue disruption using BeadBug 1.5 mm Zirconium beads (Sigma Aldrich, cat no. Z763799) in a 96-well Deep Well Plate (Fisher Scientific, cat no. 12194162) shaking at 2500rpm for 5 minutes. Genome-wide DNA methylation was profiled using the Illumina EPIC DNA methylation array (Illumina Inc), which interrogates >850,000 DNA methylation sites across the genome^69^. After stringent data quality control (see below) the BDR dataset consisted of DNA methylation estimates for 800,916 DNA methylation sites profiled in 1,221 samples (631 donors [53% male], 610 DLPFC, 611 OCC; age range = 41-104 years, median age = 84 years, mean age = 83.49 years, **Table 1**).

### Fluorescence-activated nuclei sorting (FANS) of neural cell populations from DLPFC

Neuronal-enriched, oligodendrocyte-enriched and microglia-enriched nuclei populations were isolated from ∼700mg of DLPFC tissue using a method previously described by our group^37^. First, nuclei populations were isolated from 12 donors with low neuropathology (**Table 1**) to generate reference DNA methylation profiles for purified nuclei populations for subsequent statistical deconvolution of neural cell proportions from bulk cortex DNA methylation data. Second, nuclei populations were isolated from DLPFC tissue from 15 low pathology (Braak score ≤ II) and 13 high pathology (Braak score ≥ V) BDR donors (**Table 1** and **Supplementary Table S13**) to identify cell-type-specific variable DNA methylation associated with AD pathology. Briefly, following tissue homogenization and nuclei purification using sucrose gradient centrifugation we used a FACS Aria III cell sorter (BD Biosciences) to simultaneously collect populations of NeuN+ (neuronal-enriched) and SOX10+ (oligodendrocyte-enriched) immunolabeled populations from bulk DLPFC tissue prior to genomic profiling, with the double negative fraction (microglial-enriched) and an aliquot of the ‘total’ nuclei fraction (analogous to ‘bulk’ cortex) also being collected from each tissue sample (**Supplementary Figure S19**). Nuclei suspensions were assessed for the presence of debris by adjusting the gating strategy before proceeding with nuclei capture. For each sorted population, ∼200,000 nuclei were collected for extraction of genomic DNA (**Supplementary Table S13**). Genomic DNA was isolated from each nuclei population using a standard phenol:chloroform extraction protocol^70^ and DNA methylation was profiled using the Illumina EPIC array as described above.

### DNA methylation data pre-processing and quality-control

Raw Illumina EPIC data was processed using the *wateRmelon* package as previously described^71^. Our stringent QC pipeline included the following steps: (1) checking methylated and unmethylated signal intensities and excluding poorly performing samples; (2) assessing the chemistry of the experiment by calculating a bisulphite conversion statistic for each sample, excluding samples with a conversion rate <80%; (3) identifying the fully methylated control sample included on each plate was in the correct location; (4) multidimensional scaling of sites on the X and Y chromosomes separately to confirm reported sex; (5) using the 59 SNP probes present on the Illumina EPIC array to confirm that matched samples from the same individual (but different brain regions or nuclei populations) were genetically identical and to check for sample duplications and mismatches; (6) using the *pfilter()* function in *wateRmelon* to exclude samples with >1% of probes with a detection P value□>□0.05 and probes with >1% of samples with detection P value□> □0.05; (8) using principal component (PC) analysis on data from each tissue to exclude outliers based on any of the first three PCs; and (9) the removal of cross-hybridising and SNP probes^72^. The subsequent normalization of the DNA methylation data was performed using the *dasen()* function in either *wateRmelon* or *bigmelon*^*71,73*^. The purified nuclei populations were normalized within each cell type.

### Identification of differential DNA methylation associated with neuropathology

To identify associations between variable DNA methylation and neuropathology we fitted regression models using the R (version 3.5.2) statistical environment^74^. As DNA methylation data for each donor was derived from two cortical regions, a mixed effects regression model was used, implemented with the *lme4* ^*75*^ and *lmerTest* ^*76*^ packages (see **supplementary Figure S20**). To identify DNA methylation sites associated with AD neuropathology we conducted an EWAS in which DNA methylation at each probe was regressed against the three measures of tau and amyloid pathology (Braak NFT stage, CERAD density and Thal Phase) using mixed effect regression models where age, sex, experimental batch, PC1 (which accounted for residual structure in the data) and derived neural cell proportions were included as fixed effects and individual was included as a random effect. Cell proportion estimates were derived from bulk cortex DNA methylation data using the Houseman method, implemented with minfi functions and default parameters, and a novel reference dataset generated on 12 DLPFC samples for three nuclei populations (neuronal enriched, oligodendrocyte enriched and microglial enriched) (see **Supplementary Figure S19**). Two of the three proportions (neuronal enriched and microglial enriched) were included in the model to eliminate the effects of multicollinearity. To generate *P* values, an ANOVA was conducted, comparing the full model including the three AD neuropathology measures to a null model in which the three measures were excluded. We next conducted an EWAS for each of the five neuropathology measures separately (Braak NFT stage, CERAD score, Thal Phase, Braak LB stage and TDP43-status) using the same set of covariates. Additionally, we ran analyses where cell proportions were regressed against neuropathology in each brain region using linear regression models, controlling for age and sex. To identify tissue specific effects, linear regressions models were run in each brain region for the three main AD neuropathology measures controlling for age, sex, experimental batch, PC1 and derived neural cell proportions. Finally, to further explore if there was an effect present in one cortical region and not the other we ran a heterogeneity test, where we included an interaction between neuropathology and brain region in the mixed effects models, controlling for age, sex, experimental batch, brain region, PC1, derived cell proportions and individual. EWAS results were subsequently processed using the *bacon* R package^77^, which applies a Bayesian method to adjust for inflation in EWAS.

### Meta-analysis of variable DNA methylation associated with AD pathology

Cross-cortex and DLPFC specific meta-analyses of Braak NFT stage were conducted incorporating the BDR with cohort level summary statistics from the recent meta-analysis conducted by Smith and colleagues^13^. First, the Braak NFT stage EWAS was re-run in the BDR cohort excluding 14 samples which were also present in the LBB1 cohort. In the cross-cortex meta-analysis a total of 2,939 samples (from 2,013 donors) were included (**Supplementary Table S12**). In the DLPFC meta-analysis, a total of 1,545 individuals were included (**Supplementary Table S12**). An inverse variance weighted (IVW) method was used which summarizes effect sizes from multiple independent studies by calculating the weighted mean of the effect sizes using the inverse of the variance of each study as weights. The EWAS results from each cohort were processed using the *bacon* R package^77^. A meta-analysis was then performed using the *metagen* function in the R package *meta*^*78*^, using the effect sizes and standard errors from each individual cohort to calculate weighted pooled estimates and test for significance. Probes were limited to those present in at least two of the cohorts (cross cortex n= 403,763 DNA methylation probes; DLPFC n = 402,412) and the P-value was Bonferroni corrected to control for this number of sites tested (cross cortex P <0.05/403,763 =1.24E-07; DLPFC P <0.05/402,412 = 1.24E-07). P-values are from two-sided tests and significant DMPs were taken from a fixed effects model. Pathway analyses were subsequently performed on the significant DMPs using the *methylglm* function within the *methylGSA* package developed by Ren and Kuan^79^ using the default parameters.

### Regression against AD in FANS sorted nuclei populations

To determine whether associations identified in the bulk cortex are primarily driven by alterations in specific neural cell types we used data generated on purified nuclei populations from individuals with high or low AD pathology. Briefly, we conducted an analysis of DNA methylation differences for significant sites from the bulk cortex meta-analyses comparing high and low pathology (defined as Braak high ≥ V [N = 13]; Braak low ≤ II [N = 15]) (Braak score), which was modeled as a binary variable, in the four FANS sorted nuclei populations (total nuclei [analogous to ‘bulk’ cortex], neuronal enriched, oligodendrocyte enriched and microglial enriched) separately. Linear regression models were used, whereby the significant DNA methylation sites identified in the cross-cortex and DLPFC meta-analysis were regressed against high/ low pathology status controlling for age, sex, and batch (brain bank). The results were then compared to the meta-analysis results where a binomial test (sign test) was used to statistically evaluate consistency in direction of effect across the analyses.

## Data availability

BDR DNA methylation data are available via the Dementias Platform UK (DPUK) data portal (https://portal.dementiasplatform.uk/) and the Gene Expression Omnibus (GEO) at accession number GSE197305. Analysis scripts used in this manuscript are available on GitHub (https://github.com/gemmashireby/BDR_neuropathology_EWAS).

## ACKNOWLEDGEMENTS

G.S. was supported by a PhD studentship from the Alzheimer’s Society. E.H., E.D. and J.M. were supported by the Medical Research Council (MRC) grant K013807. Data analysis was undertaken using high-performance computing supported by a Medical Research Council (MRC) Clinical Infrastructure award to J.M. (M008924). The analysis of FANS-purified nuclei was supported by Alzheimer’s Research UK (ARUK) grant ARUK-PPG2018A-010 to E.D. DNA methylation data generated in the Brains for Dementia Research cohort was supported by the Alzheimer’s Society and Alzheimer’s Research UK (ARUK). The BDR is jointly funded by Alzheimer’s Research UK (ARUK) and the Alzheimer’s Society in association with the Medical Research Council. The South West Dementia Brain Bank is part of the Brains for Dementia Research program, jointly funded by Alzheimer’s Research UK (ARUK) and Alzheimer’s Society, and is also supported by BRACE (Bristol Research into Alzheimer’s and Care of the Elderly) and the Medical Research Council (MRC).

## TABLE LEGEND

**Table 1: Characteristics of the samples profiled in this study. IQR = interquartile range. SD = standard deviation.**

