## Supplementary Figures for "DNA methylation signatures of Alzheimer’s disease neuropathology in the cortex are primarily driven by variation in non-neuronal cell-types"

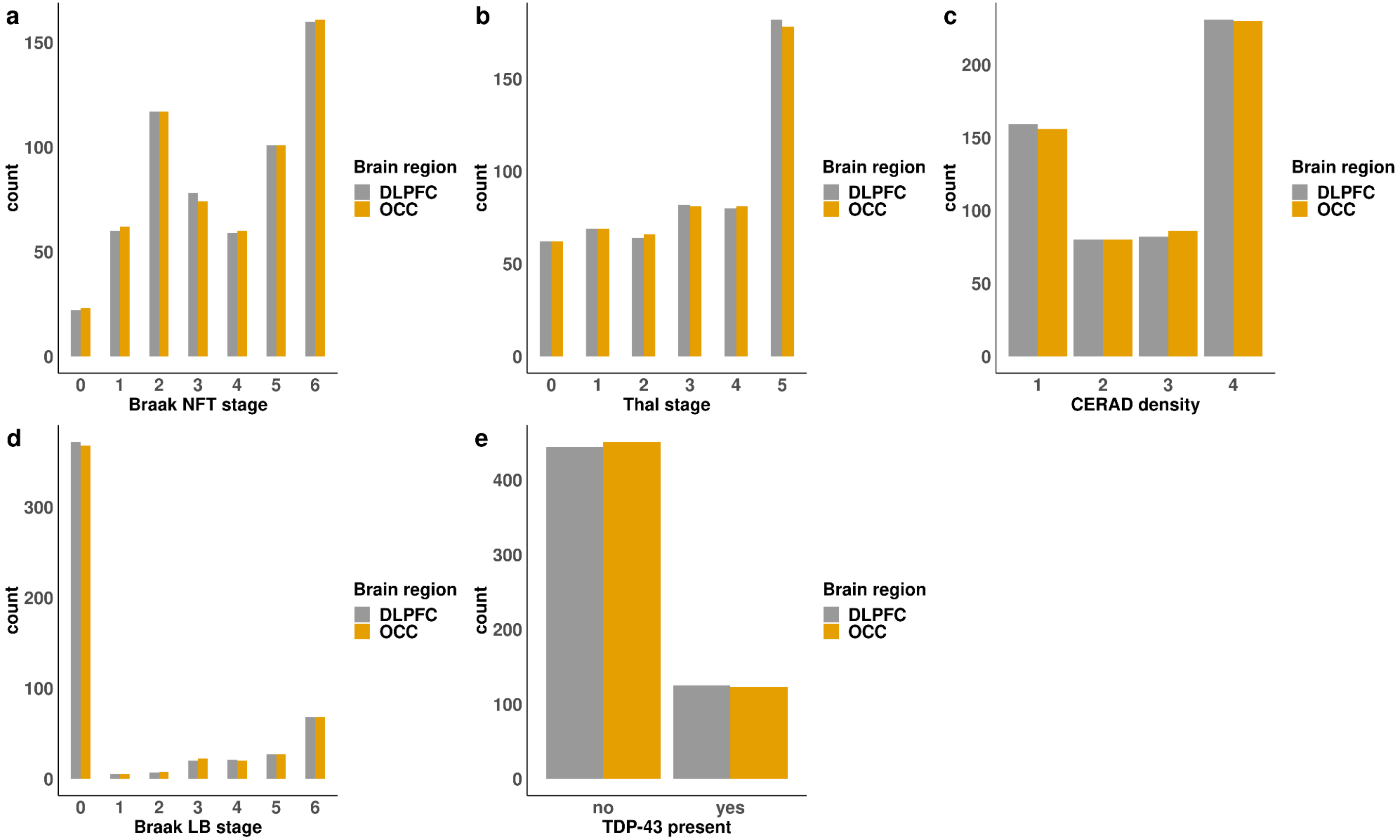


**Figure S1: Levels of neuropathology across BDR donors included in this study.** Shown is the distribution of neuropathology as measured by **a)** Braak NFT stage, **b)** Thal stage, **c)** CERAD density, **d)** Braak LB stage and **e)** TDP43 status.

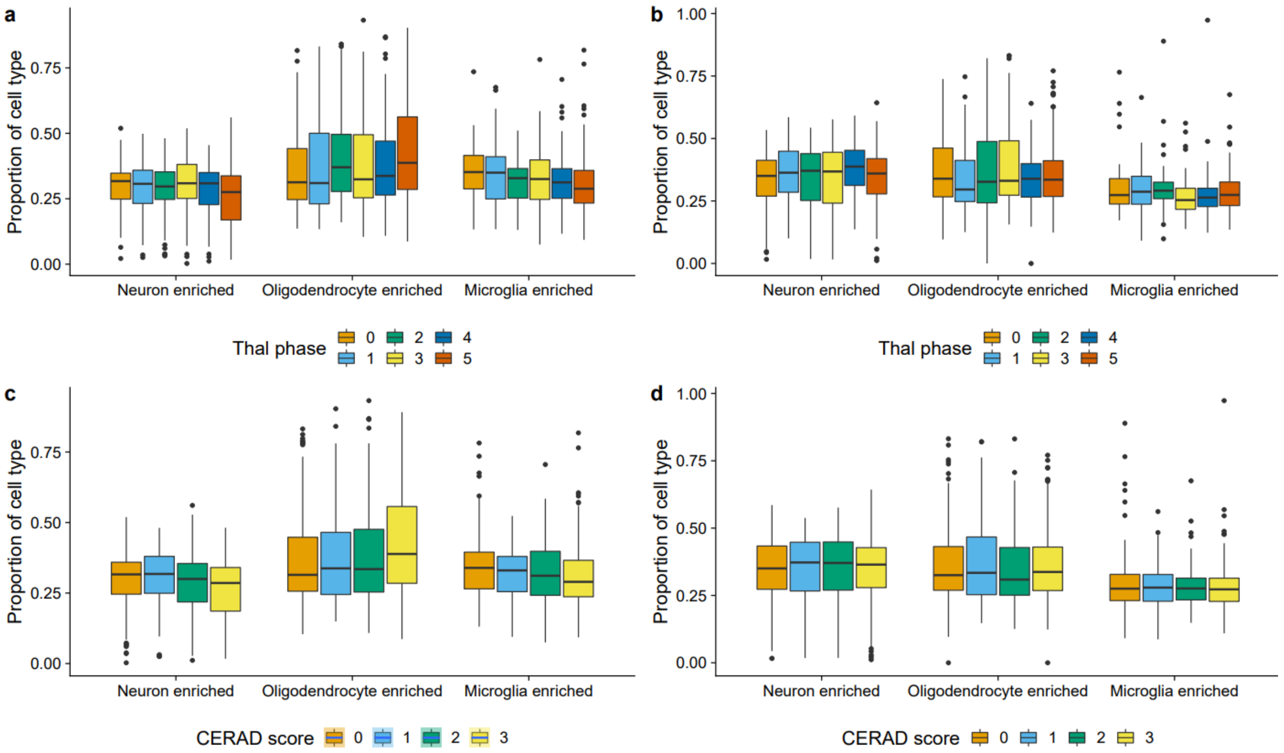


**Figure S2: Elevated amyloid pathology is associated with cell proportion estimates derived from DNA methylation data in the DLPFC but not the OCC. a)** Thal phase was significantly associated with the proportion of neurons (effect size = -1.38, P = 0.006), oligodendrocytes (effect size = 1.26, P = 0.003) and microglia (effect size = -2.39, P = 0.001) in the DLPFC using neural cell proportion estimates derived from ‘bulk’ DNA methylation data. Boxplots plots for the estimated proportion of cell type across Braak NFT stages are shown, where the box in the middle represents the interquartile range (IQR), and the whisker lines represent the minimum (quartile 1 – 1.5 x IQR) and the maximum (quartile 3 + 1.5 x IQR). Amyloid pathology is shown on the x-axis split by cell-type and estimated cell proportions are shown on the y-axis. **b)** In contrast no associations between Thal phase and cell proportion estimates derived from DNA methylation data were observed in the OCC (P > 0.008). **c)** CERAD score was significantly associated with the proportion of neurons (effect size = -1.38, P = 0.006), oligodendrocytes (effect size = 0.970, P = 0.001) and microglia (effect size = -1.39, P = 0.004) in the DLPFC using neural cell proportion estimates derived from ‘bulk’ DNA methylation data. **d)** In contrast no associations between CERAD score and cell proportion estimates derived from DNA methylation data were observed in the OCC (P > 0.008).

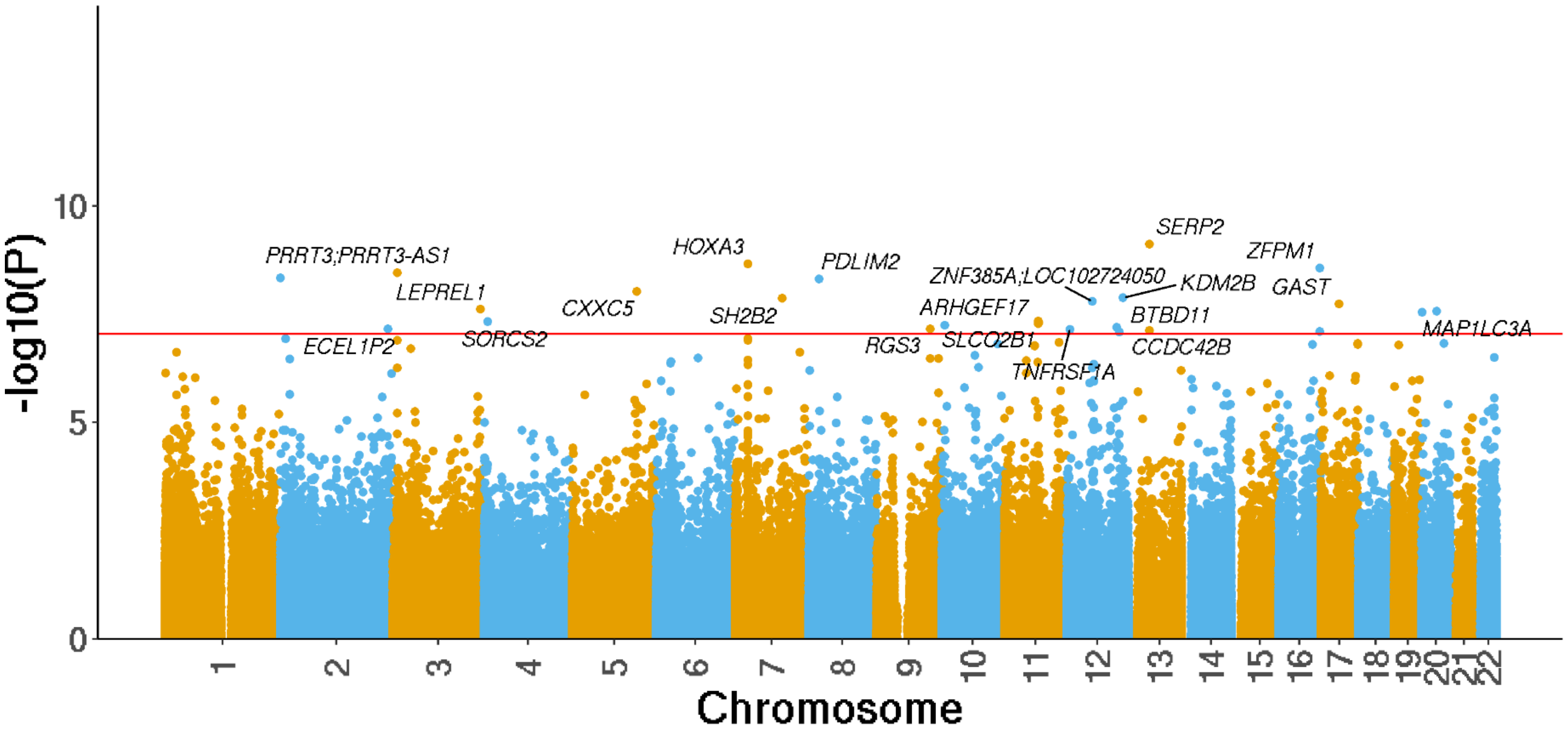


**Figure S3: Manhattan plot highlighting cortical DMPs significantly associated with Braak neurofibrillary tangle stage.** Genes annotated to significant DMPs are labelled. The x-axis shows chromosomes 1-22 and the y-axis shows -log10(P), with the horizontal red line representing experiment wide significance (P < 9e-8). A complete list of results is given in **Supplementary Table S3.**

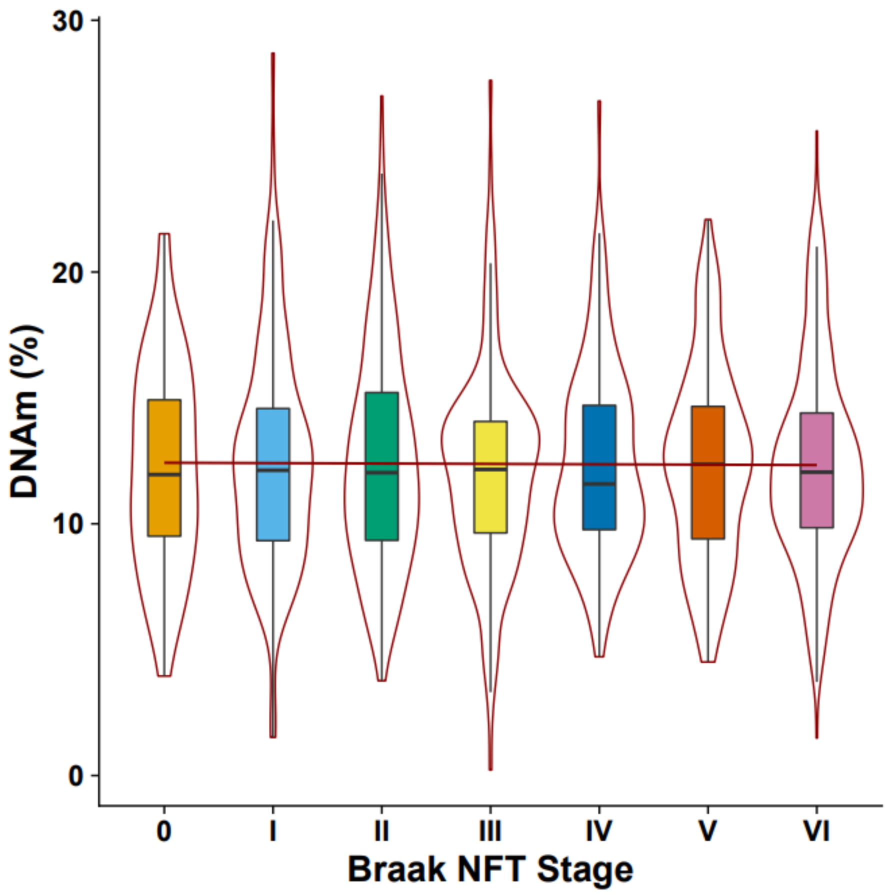


**Figure S4: The top-ranked DMP associated with tau pathology.** Shown is mean DNA methylation levels (adjusted for covariates, see **Methods**) at cg16021126 (annotated to SERP2) across Braak NFT stages with significant hypermethylation with elevated pathology (P = 7.48e-10, effect size = 0.286%). The middle box represents the interquartile range (IQR) and the whisker lines represent the minimum (quartile 1 – 1.5 x IQR) and the maximum (quartile 3 + 1.5 x IQR). Pathology stage (Braak NFT) is shown on the x-axis with DNA methylation level, adjusted for covariates, shown on the y-axis.

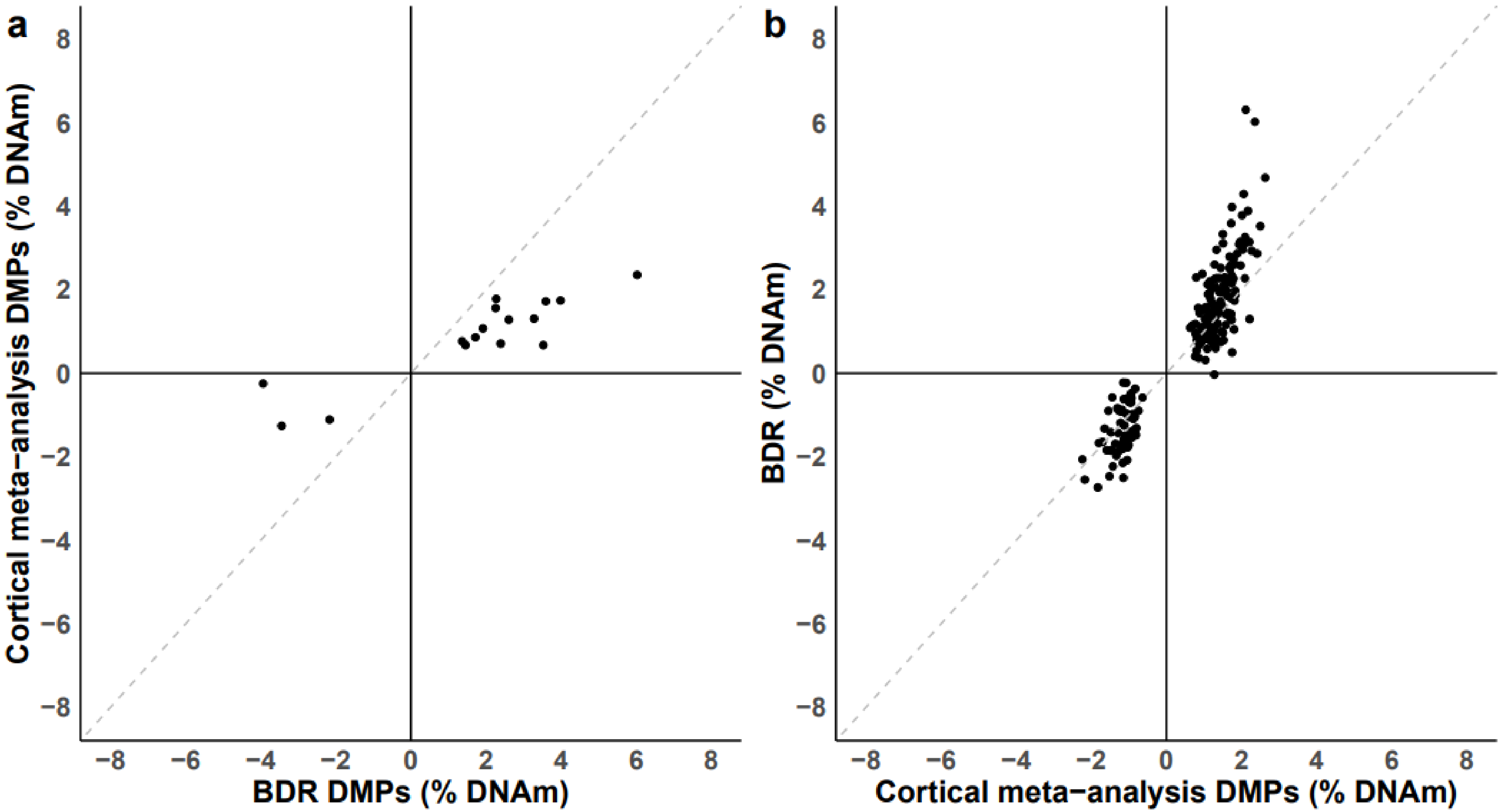


**Figure S5: Effects sizes at DNA methylation sites associated with Braak neurofibrillary tangle (NFT) stage in BDR are highly consistent with those reported in a recent meta-analysis of Braak NFT stage[13]**. Shown for sites tested in both analyses is a comparison of **a)** effect sizes at Braak NFT-associated DMPs identified in the BDR cohort (n = 16) with those from the recent meta-analysis of Braak NFT stage[13] (direction of effect = 100% concordant, sign test P =1.53e-05), **b)** effect sizes at Braak NFT-associated DMPs identified in the recent meta-analysis (n = 208) with those from the BDR cohort (direction of effect = 100% concordant, sign test P =5.08e-61) Concordant = percentage of DNA methylation sites which have the same direction of effect. Grey dashed line represents y=x.

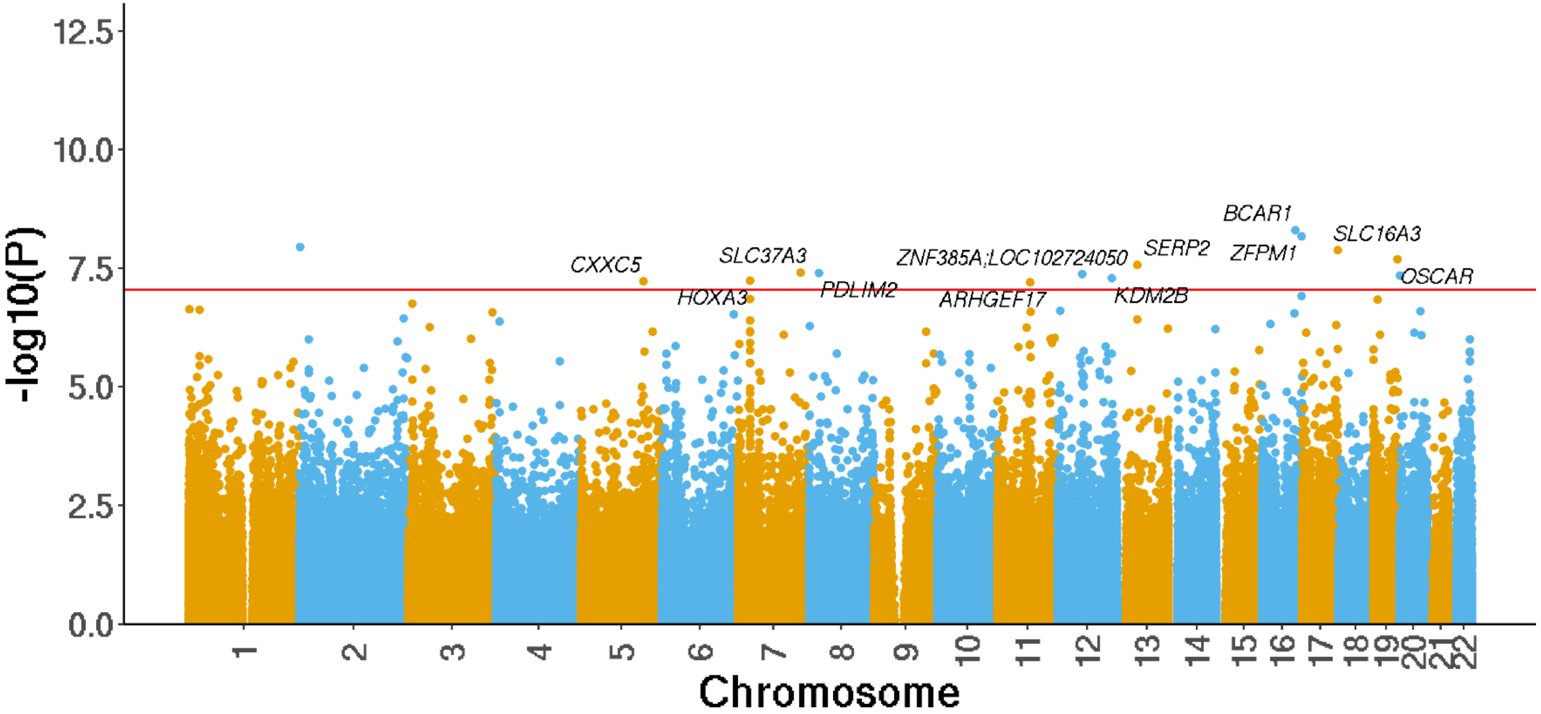


**Figure S6: Manhattan plot highlighting significant cortical DMPs associated with CERAD density. Genes** annotated to significant DMPs are labelled. The x-axis shows chromosomes 1-22 and the y-axis shows -log10(P), with the horizontal red line representing experiment wide significance (P < 9e-8). A complete list of results is given in **Supplementary Table S3.**

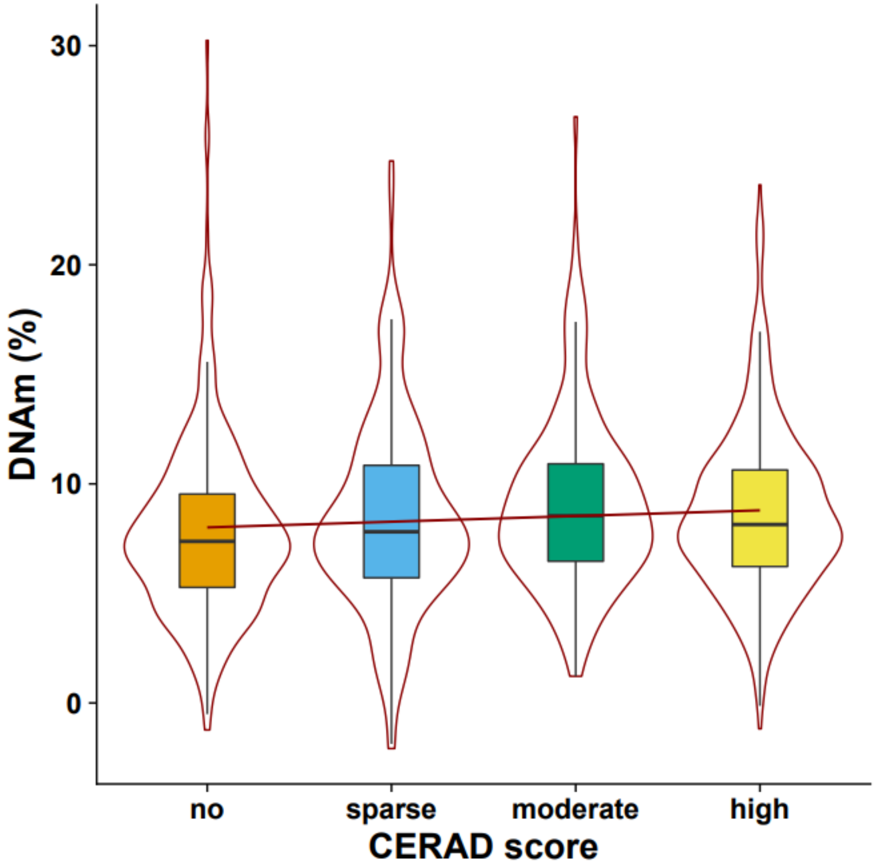


**Figure S7: The top-ranked DMP associated with the CERAD score measure of amyloid pathology.** The top ranked DMP associated with CERAD score was cg13515047 annotated to *BCAR1*, which was significantly hypermethylated with elevated pathology (P = 4.96e-09, effect size = 0.442%). Violin plots for the DNA methylation values (adjusted for covariates, see **Methods**) across CERAD score are shown, where the box in the middle represents the interquartile range (IQR), and the whisker lines represent the minimum (quartile 1 – 1.5 x IQR) and the maximum (quartile 3 + 1.5 x IQR). Pathology stage (CERAD) is shown on the x-axis with DNA methylation level, adjusted for covariates, shown on the y-axis.

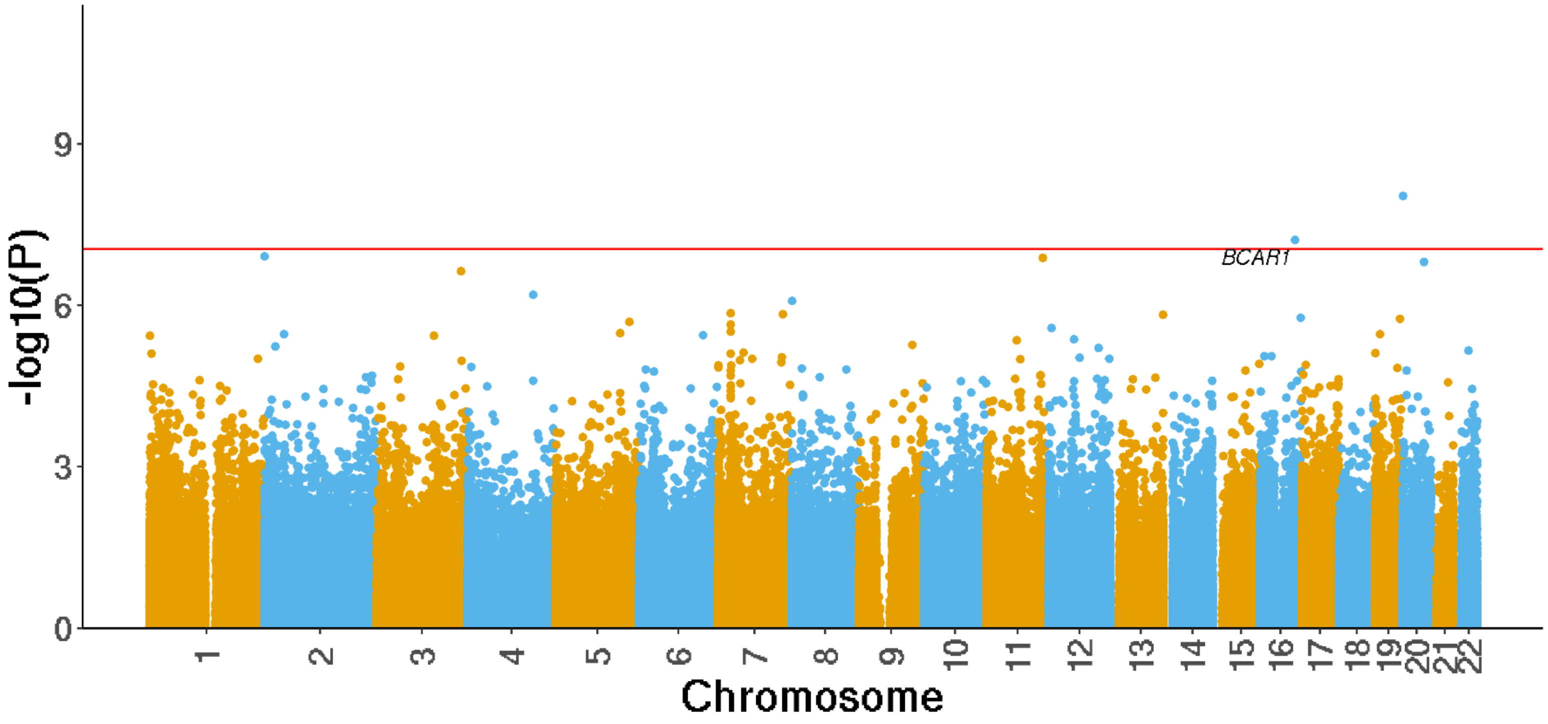


**Figure S8: Cortex EWAS of Thal Phase.** Genes annotated to significant DMPs are labelled. The x-axis shows chromosomes 1-22 and the y-axis shows -log10(P), with the horizontal red line representing experiment wide significance (P < 9e-8). A complete list of results is given in **Supplementary Table S3.**

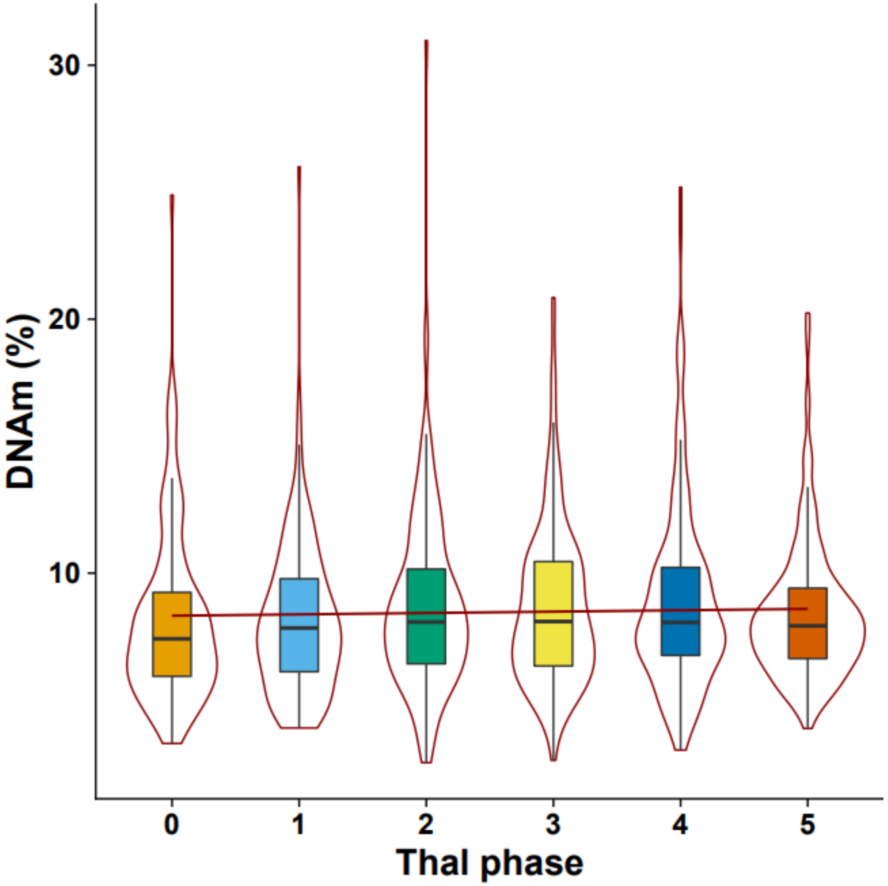


**Figure 9: The top-ranked DMP associated with the Thal phase measure of amyloid pathology.** The top ranked DMP associated with Thal phase was cg11658414 (not annotated to a gene), which was significantly hypermethylated with elevated pathology (P = 9.11E-09, effect size = 0.299%). Violin plots for the level of DNA methylation (adjusted for covariates, see **Methods**) across Thal groups are shown, where the box in the middle represents the interquartile range (IQR), and the whisker lines represent the minimum (quartile 1 – 1.5 x IQR) and the maximum (quartile 3 + 1.5 x IQR). Pathology stage (Thal) is shown on the x-axis with DNA methylation level, adjusted for covariates, shown on the y-axis.

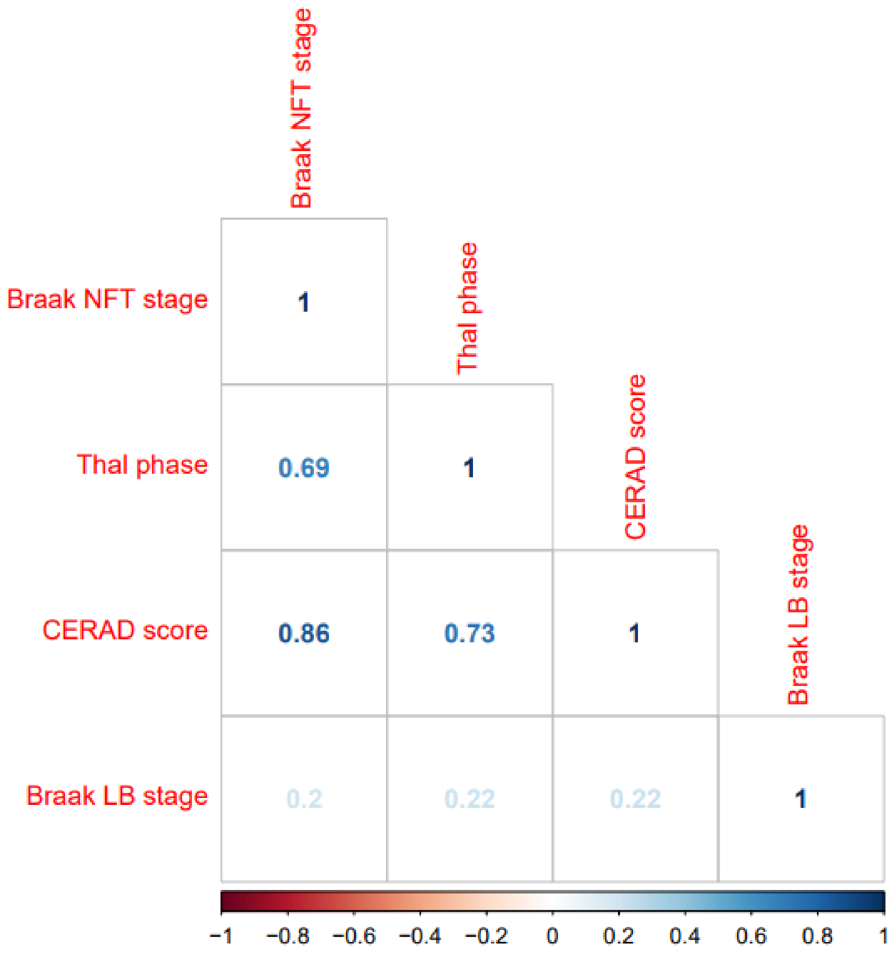


**Figure S10: Heatmap showing a strong correlation between different measures of AD neuropathology across samples included in this study.** There is a weaker correlation between measures of AD pathology and Braak LB stage. NFT- neurofibrillary tangles, LB – Lewy body.

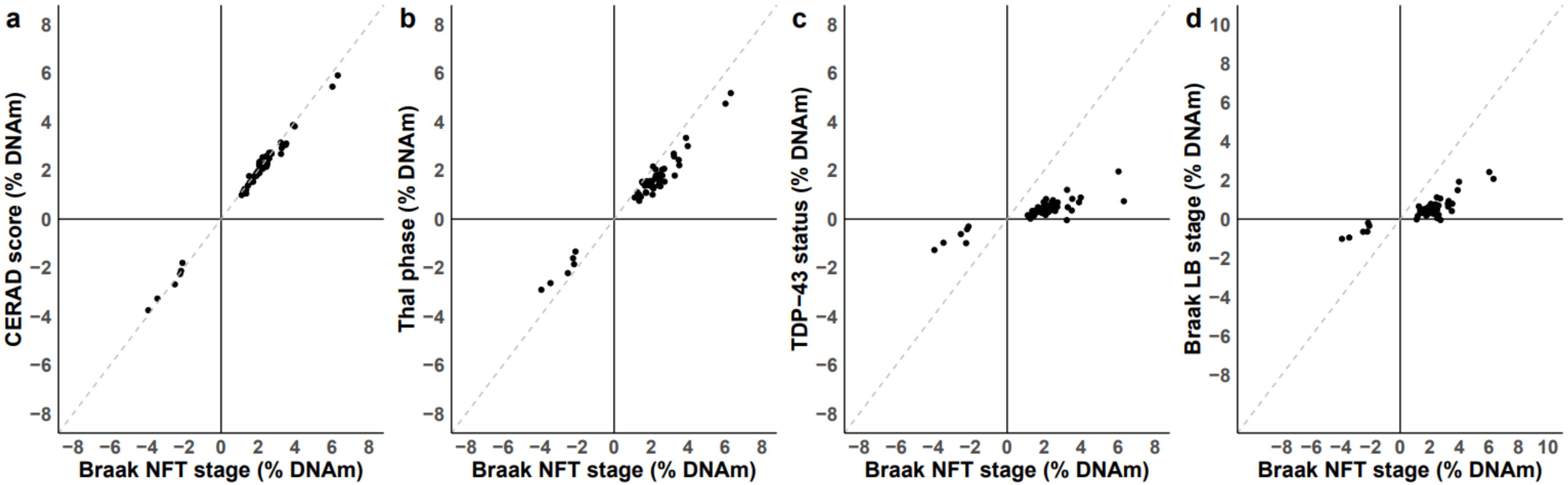


**Figure S11: Effects sizes at DNA methylation sites associated with Braak neurofibrillary tangle (NFT) are highly consistent with those from EWAS analyses of other dementia neuropathology measures.** Shown are the effect sizes for 26 tau-associated DMPs identified in the BDR cohort comparing the results from an EWAS of Braak NFT stage with **a)** CERAD score (direction of effect = 100% concordant, sign test P =1.39e-17), **b)** Thal Phase (direction of effect = 100% concordant, sign test P = 1.39e-17), **c)** TDP-43 status (direction of effect = 98% concordant, sign test P = 7.91e-16) and **d)** Braak LB Stage (direction of effect = 96% concordant, sign test P =2.22e-14). Grey dashed line represents y=x.

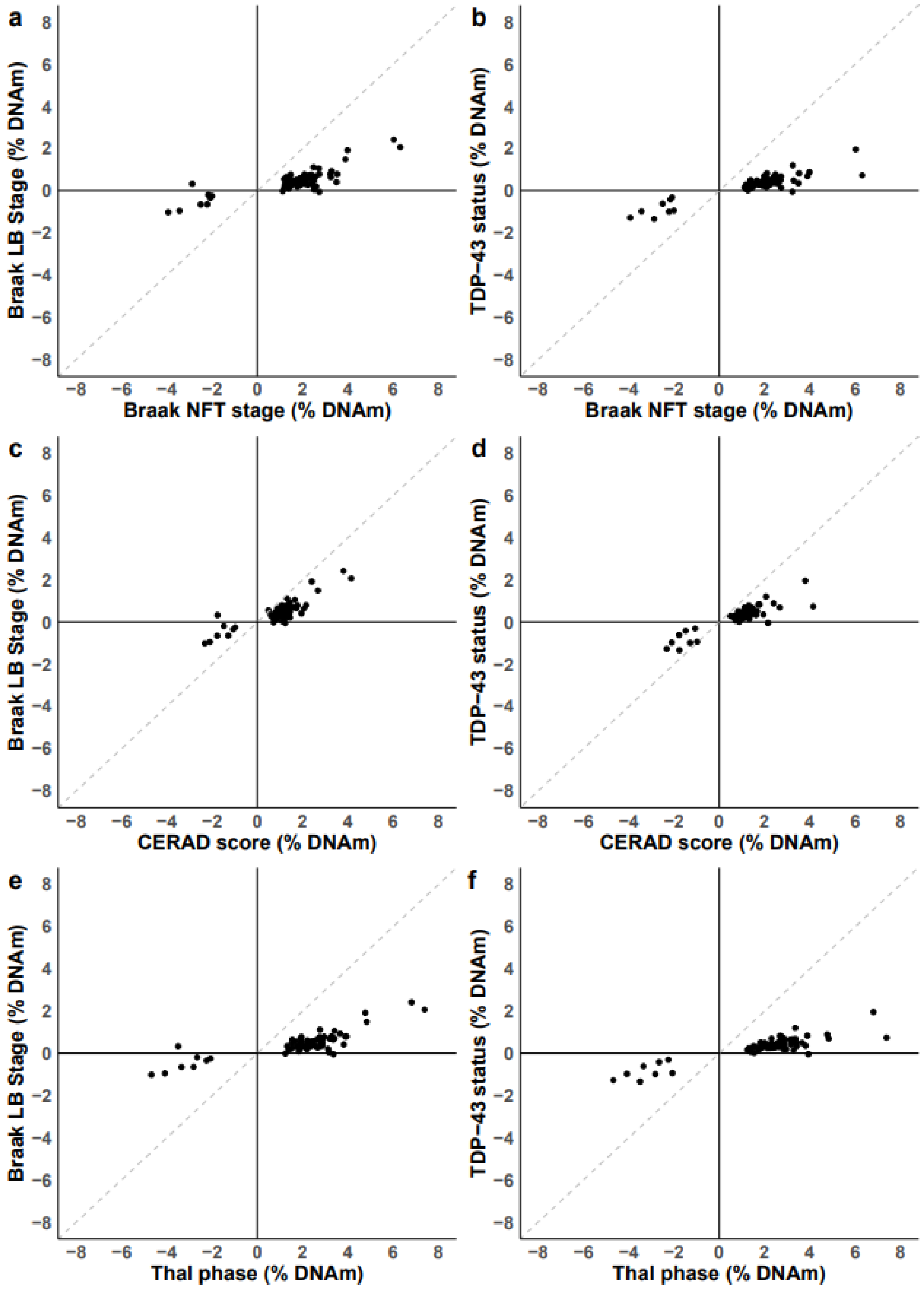


**Figure S12: Effects sizes at the 67 cortical DNA methylation sites associated with AD neuropathology are highly consistent across EWAS analyses of individual dementia neuropathology measures.** Shown are the effects sizes for the 67 core-AD neuropathology DNA methylation sites from EWAS of the separate neuropathology measures comparing **a)** Braak NFT stage and Braak LB stage (96% concordant; sign test p-value= 3.4e-16), **b)** Braak NFT stage and TDP-43 status (99% concordant; sign test p-value=4.16e-19), **c)** Thal phase and Braak LB Stage (96% concordant; sign test p-value=3.4e-16), **d)** Thal phase and TDP-43 status (99% concordant; sign test p-value= 4.16e-19), **e)** CERAD score and Braak LB Stage (96% concordant; sign test p-value= 3.4e-16), and **f)** the CERAD score and TDP-43 status EWAS (99% concordant; sign test p-value= 4.16e-19). Grey dashed line represents y=x.

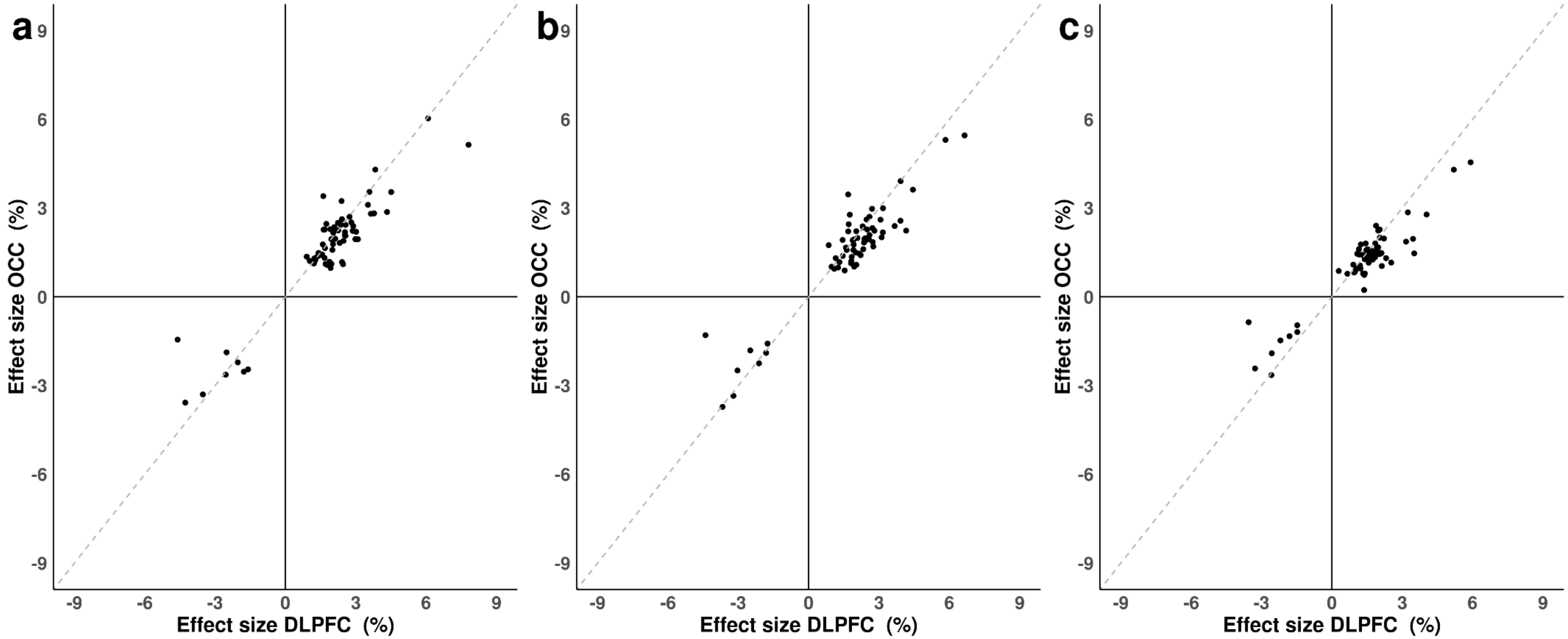


**Figure S13: Effects sizes at the 67 cortical DNA methylation sites associated with AD neuropathology are highly consistent across both cortical regions.** Shown are the effects sizes for the 67 core-AD DNA methylation sites from EWAS of each AD neuropathology measure conducted in the DLPFC and OCC separately where **a)** Braak NFT stage (100% concordant; sign test P =6.78e-21**), b)** CERAD score (100% concordant; sign test *P* =6.78e-21), and **c)** Thal phase (100% concordant; sign test P =6.78e-21). Grey dashed line represents y=x. OCC=occipital cortex; DLPFC = dorsolateral prefrontal cortex.

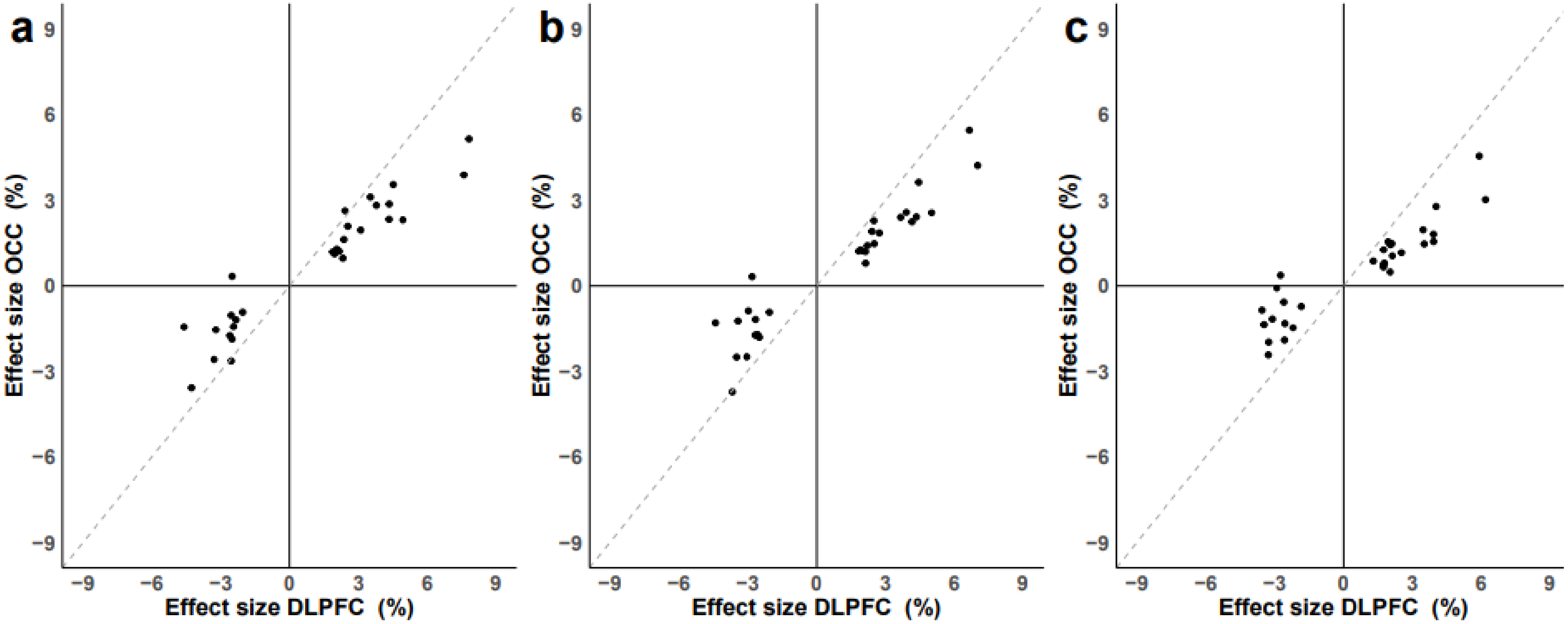


**Figure S14: Effect sizes at DNA methylation sites associated with AD neuropathology in the DLPFC are consistent with those identified in the OCC.** Shown are the effects sizes for the 30 AD-associated DMPs identified in the DLPFC between the two cortical regions from EWAS analyses of **a)** Braak NFT stage (direction of effect = 97% concordant, sign test P =2.89e-08), **b)** CERAD score (direction of effect = 97% concordant, sign test P =2.89e-08), and **c)** Thal phase (direction of effect = 97% concordant, sign test P =2.89e-08). Grey dashed line represents y=x. OCC=occipital cortex; DLPFC = dorsolateral prefrontal cortex.

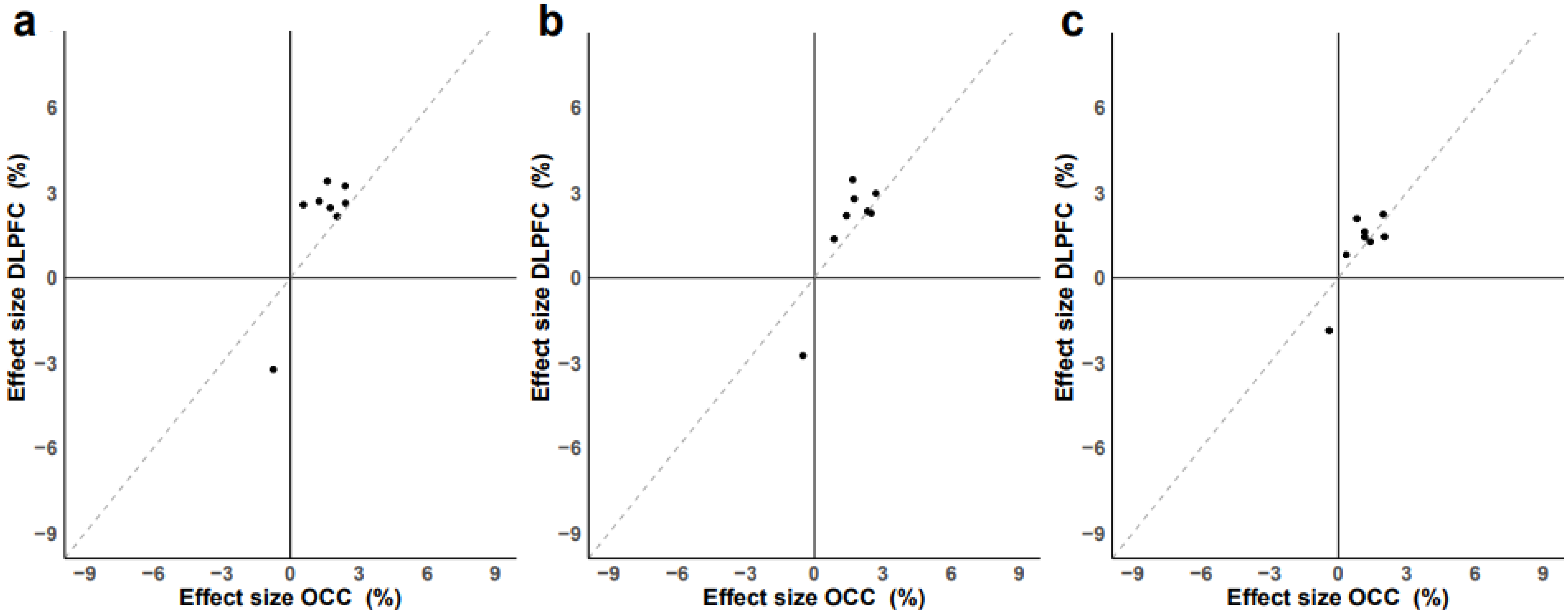


**Figure S15: Effect sizes at DNA methylation sites associated with AD neuropathology in the OCC are consistent with those identified in the DLPFC.** Shown are the effect sizes at the 8 AD-associated DMPs identified in the OCC between the two cortical regions from EWAS analyses of **a)** Braak NFT stage (direction of effect = 100% concordant, sign test P =0.00391), **b)** CERAD score (direction of effect = 100% concordant, sign test P =0.00391), and **c)** Thal phase (direction of effect = 100% concordant, sign test P =0.00391). Grey dashed line represents y=x. OCC=occipital cortex; DLPFC = dorsolateral prefrontal cortex.

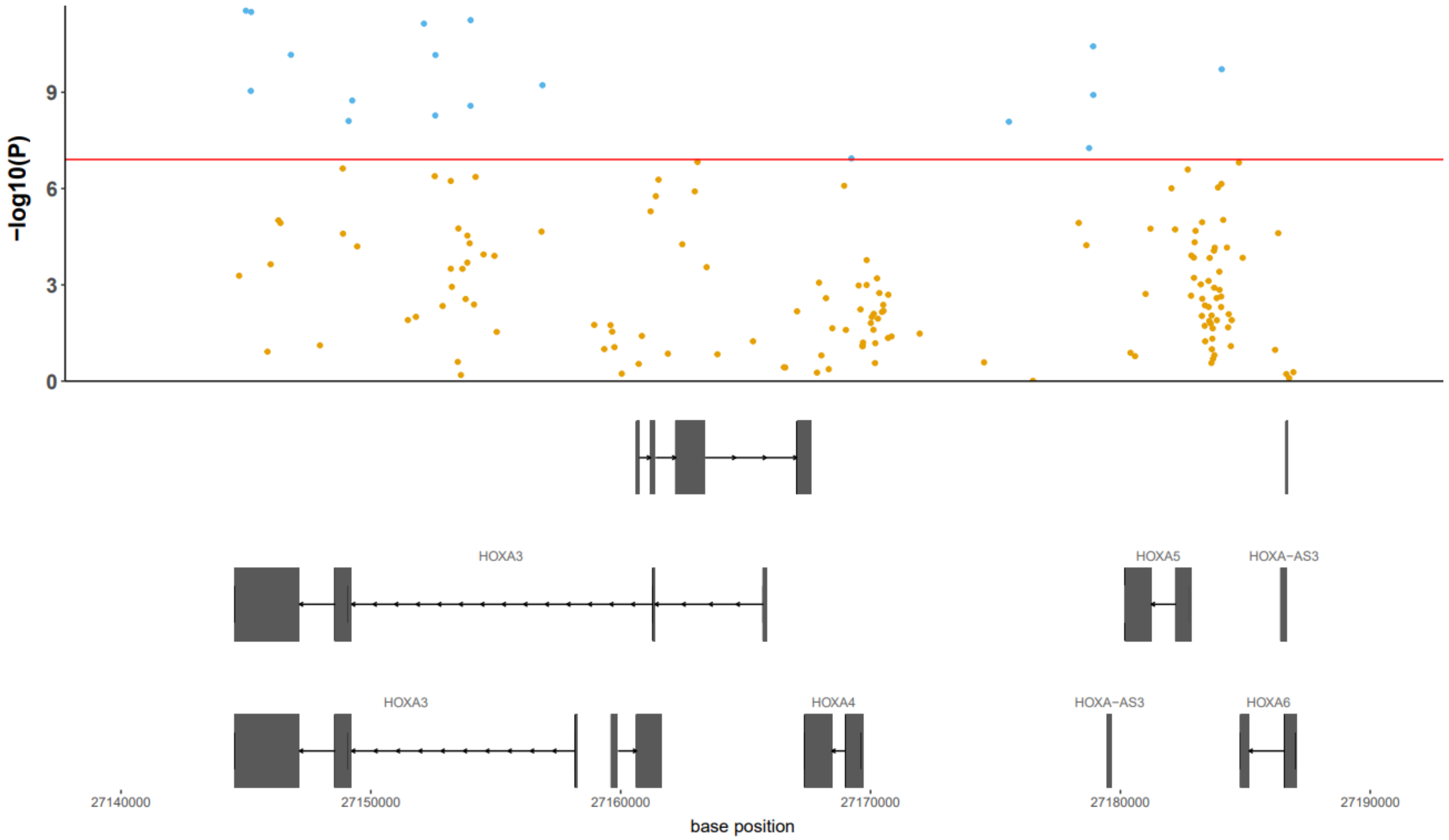


**Figure S16: Multiple DMPs in the HOXA region were associated with tau pathology in the cortex meta-analysis.** Shown in the top panel is a zoomed in Manhattan plot around the HOXA region on chromosome *7*, where the x-axis represents the base position, and the y-axis represents the –log10 P-value, with each point on the plot representing a DNA methylation site. The red horizontal line represents experiment wide significance ($P < 1.24e-07$). The gene track shows the locations of the genes within in this region (hg19).

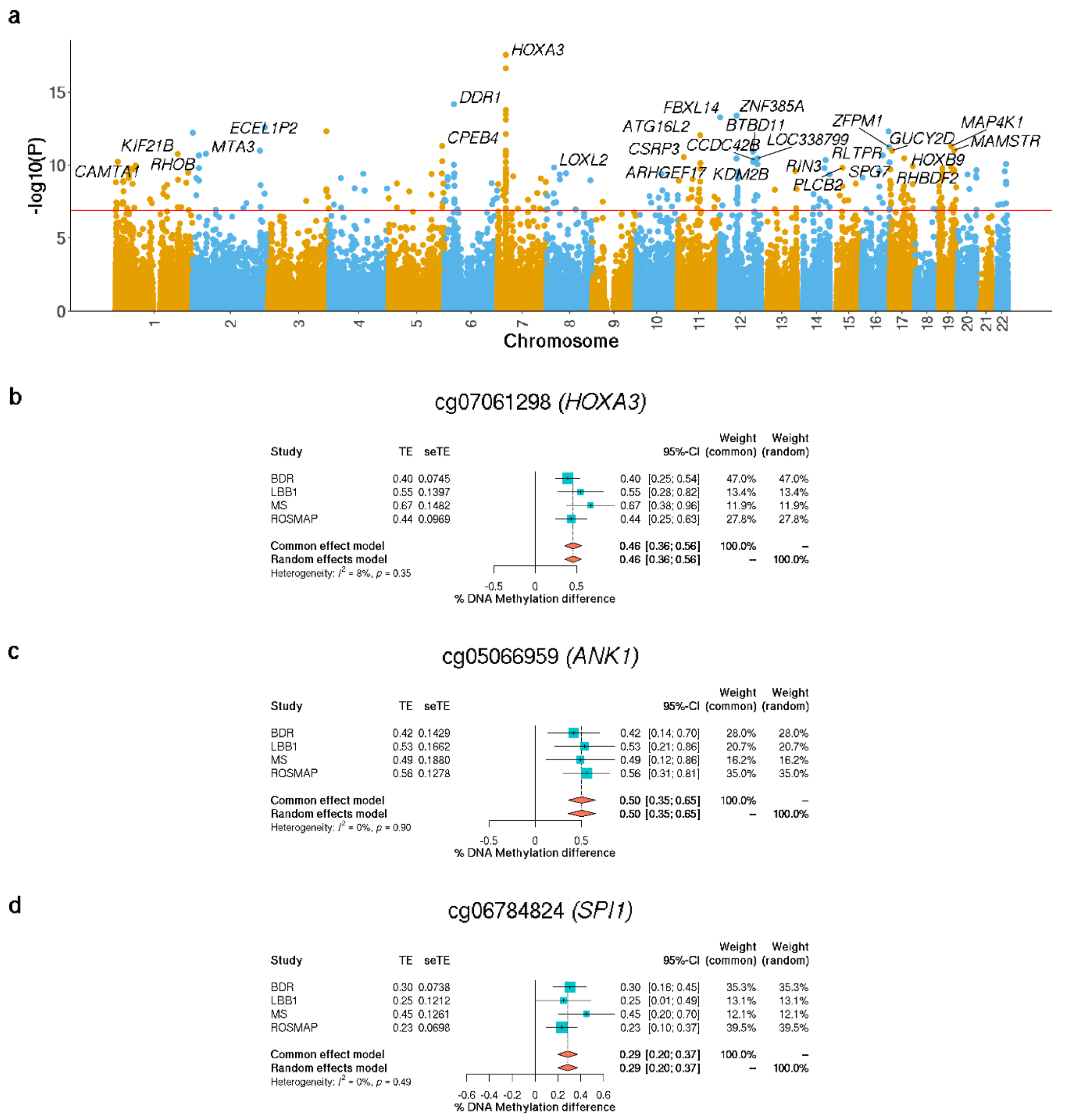


**Figure S17: Differentially methylated positions identified in the DLPFC meta-analysis are annotated to genes strongly implicated in Alzheimer's disease. a)** Manhattan plot highlighting significant DMPs associated with Braak NFT stage in a cross-cohort meta-analysis. Genes annotated to the 50 most significant DMPs are labelled. The x-axis shows chromosomes 1-22 and the y-axis shows -log10(P), with the horizontal red line representing experiment wide significance (P < 9E-8). A complete list of results is given in **Supplementary Table S12. b)** Across all cohorts Braak NFT Stage is associated with hypermethylation at cg22962123 (P = 2.31E-17) which is annotated to *HOXA3*, a gene previously implicated in EWAS of AD[13,45,46]. **c)** Across all studies Braak NFT Stage is associated with hypermethylation at cg05066959 (P = 4.36E-10) which is annotated to *ANK1*, a gene previously implicated in EWAS of AD. **d)** Across all studies Braak NFT Stage is associated with hypermethylation at cg06784824 (P = 8.97E-10) which is annotated to *SPI1*, a gene previously implicated in GWAS[7,54] and EWAS[13] of AD. The X-axis shows the effect size (% DNA methylation difference per SD increase in Braak NFT stage), with squares representing effect size and arms indicating the 95% confidence intervals.

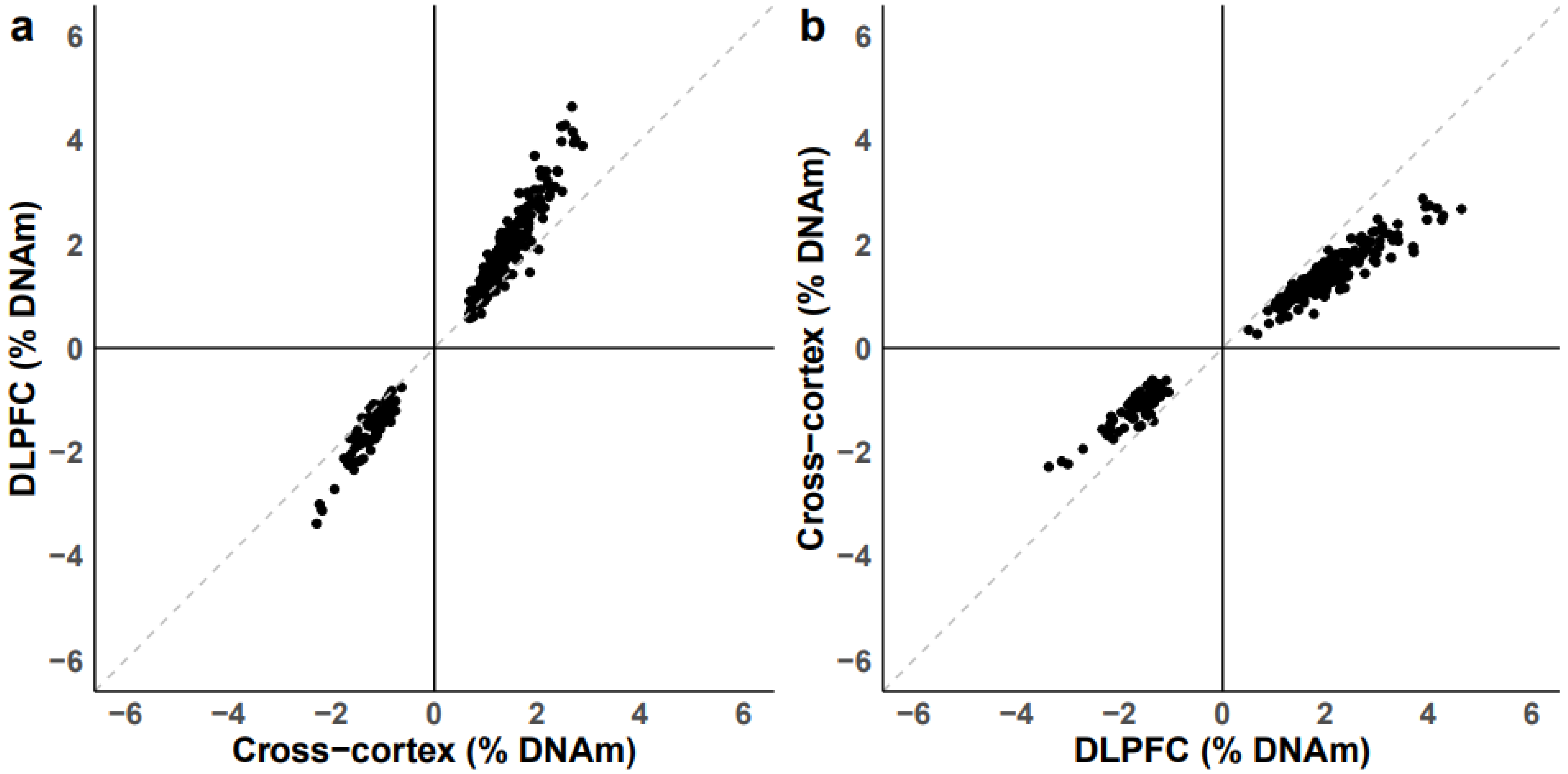


**Figure S18: Effects sizes from EWAS meta-analyses of Braak neurofibrillary tangle (NFT) stage undertaken in the DLPFC are highly consistent with those from a cross-cortex meta-analysis. a)** Shown are the effect sizes at 334 DMPs identified in the cross-cortex meta-analysis to those same DNA methylation sites in the DLPFC meta-analysis (direction of effect = 100% concordant, sign test P = 2.86E-101). **b)** Shown are the effect sizes at 300 DMPs identified in the DLPFC meta-analysis compared to those same DNA methylation sites in the cross-cortex meta-analysis (concordant = 100%, sign test P = 4.91E-91). Grey dashed line represents y=x.

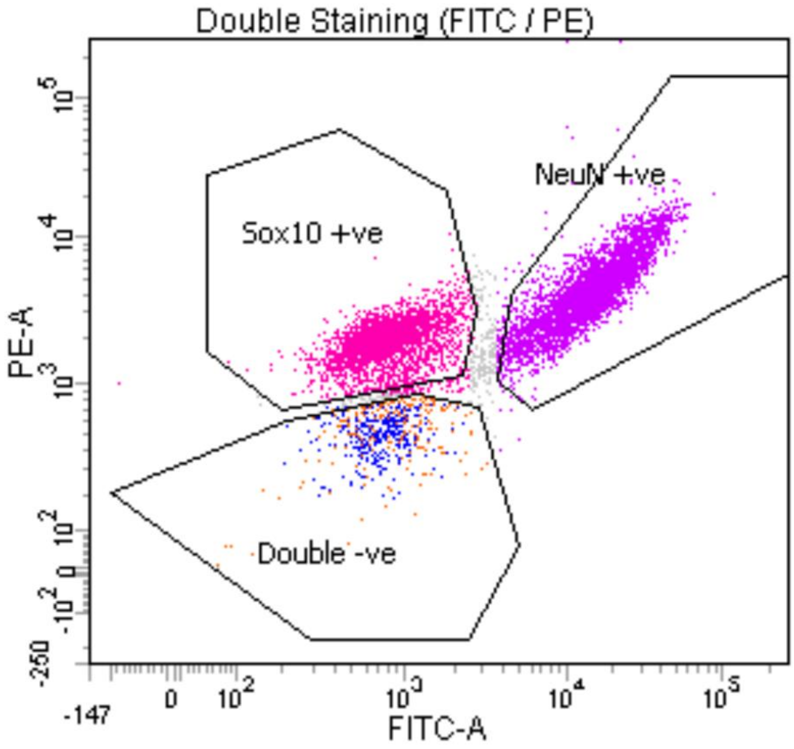


**Figure S19: Isolation of cell-type-enriched nuclei populations from DLPFC tissue using fluorescence activated nuclei sorting (FANS).** Shown is a representative example from one BDR DLPFC sample highlighting the FANS separation of three discrete nuclei populations. Purple = NeuN+ (neuronal enriched), pink=SOX10+ (oligodendrocyte enriched), blue = double negative (microglial enriched). IRF8+, a specific microglial marker, is shown in orange highlighting the overlap with the double negative population.

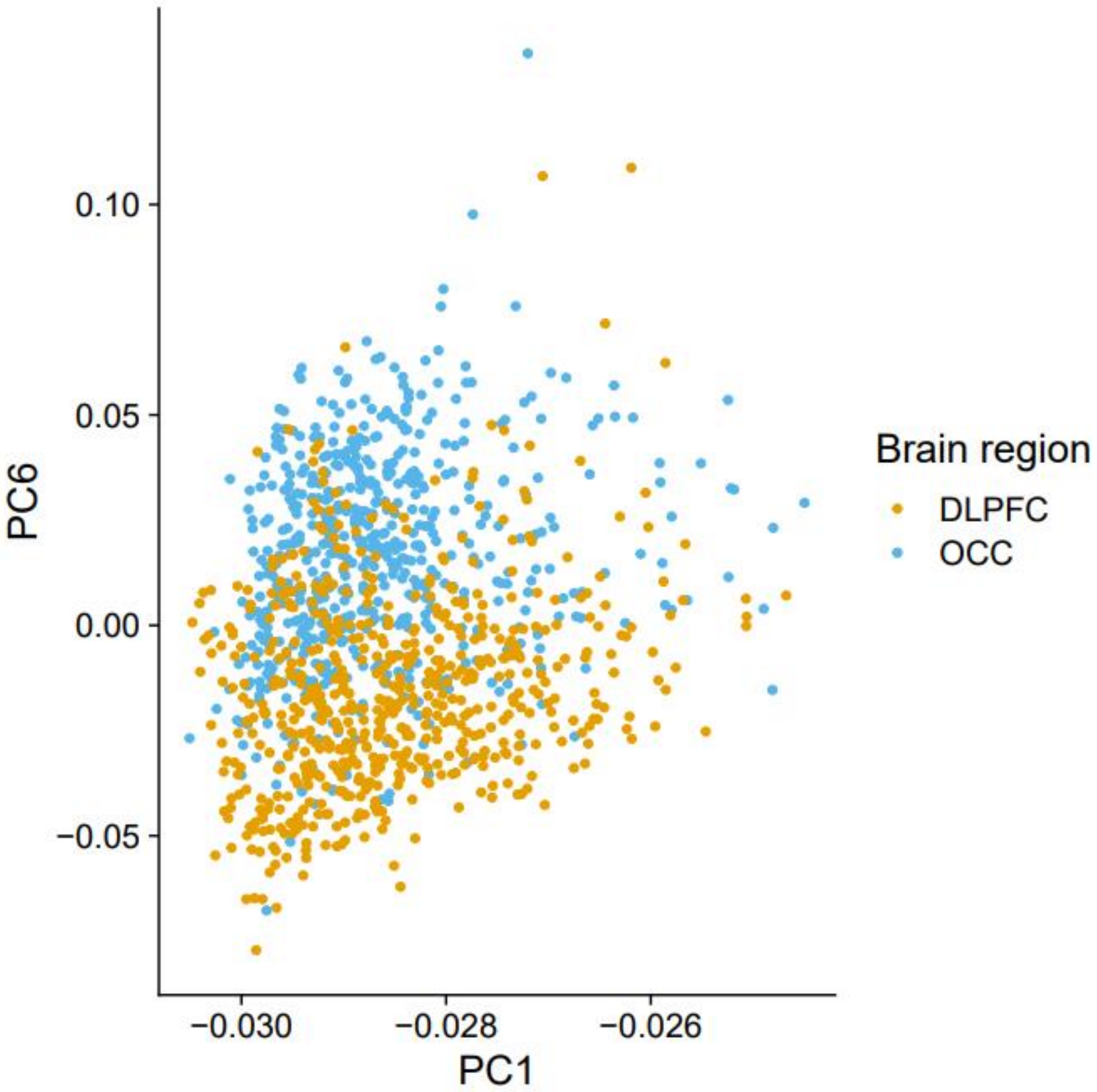


**Figure S20: Principal components of DNA methylation data from both cortical regions largely overlap.** PC6 is most correlated with brain region (r=-0.54; P = 5.4e-92) but only explains *0.08%* of the variance.
